# Movie-watching outperforms rest for functional connectivity-based prediction of behavior

**DOI:** 10.1101/2020.08.23.263723

**Authors:** Emily S. Finn, Peter A. Bandettini

**Affiliations:** Section on Functional Imaging Methods, Laboratory of Brain and Cognition, National Institute of Mental Health, Bethesda, Md. USA; Department of Psychological and Brain Sciences, Dartmouth College, Hanover, N.H. USA

## Abstract

A major goal of human neuroscience is to relate differences in brain function to differences in behavior across people. Recent work has established that whole-brain functional connectivity patterns are relatively stable within individuals and unique across individuals, and that features of these patterns predict various traits. However, while functional connectivity is most often measured at rest, certain tasks may enhance individual signals and improve sensitivity to behavioral differences. Here, we show that compared to the resting state, functional connectivity measured during naturalistic viewing—i.e., movie watching—yields more accurate predictions of trait-like phenotypes in the both cognitive and emotional domains. Traits could be predicted using less than three minutes of data from single video clips, and clips with highly social content gave the most accurate predictions. Results suggest that naturalistic stimuli amplify individual differences in behaviorally relevant brain networks.

## Introduction

Individual patterns of whole-brain functional connectivity are stable and unique enough to serve as a “fingerprint” that can identify people across time and brain states. Features of these patterns predict a growing list of phenotypes, including fluid intelligence (Finn et al., 2015), sustained attention (Rosenberg et al., 2016), and personality traits (Hsu et al., 2018), among others. Characterizing these patterns could give insight into the systems-level structure of trait-like phenotypes, and refining such predictive models may lead to biomarkers of present or future health status and other outcomes (Gabrieli et al., 2015; Dubois and Adolphs, 2016; Finn and Constable, 2016; Woo et al., 2017).

Yet despite its trait-like aspects, functional connectivity also shows a considerable state-like component such that task demands modulate connectivity patterns at both the group and individual level (Waites et al., 2005; Finn et al., 2017; Gratton et al., 2018; Rosenberg et al., 2020). These observations raise the question, what is the best brain state is for studying individual differences? Investigators often default to rest as a supposedly neutral backdrop, and indeed, resting-state acquisitions have several advantages: they are easy to acquire and standardize across sites and populations, less vulnerable to performance or motivation confounds, and relatively robust to practice or repetition effects. However, rest is also more susceptible to arousal confounds and can reduce subject compliance, especially in hard-to-scan populations (Vanderwal et al., 2015; Huijbers et al., 2017). Other candidate states include the myriad tasks traditionally used in cognitive and social psychology, or naturalistic paradigms, in which participants watch movies or listen to stories. The latter are growing in popularity as a window into brain activity under rich and engaging conditions that are more ecologically valid than highly controlled tasks (Sonkusare et al., 2019).

Early work using movies and other naturalistic stimuli focused on how they tend to synchronize brain responses across people, resulting in similar spatiotemporal activity patterns in individuals experiencing the same stimulus (Hasson et al., 2004; Nastase et al., 2019). One might therefore expect that these conditions would quench between-subject variability, making them undesirable for studying individual differences. But in fact, observable individual differences persist atop this shared response, in both activity (Finn et al., 2020) and functional connectivity (Geerligs et al., 2015; Vanderwal et al., 2017; Wang et al., 2017). These findings complement the somewhat paradoxical observation that even among traditional paradigms (e.g., n-back, finger tapping, emotional faces), conditions that make subjects’ connectivity profiles more similar to one another also make them easier to identify (Finn et al., 2017). One interpretation of these findings is that any task, but especially rich, engaging tasks, constrain the functional connectivity space in a way that reduces overall between-subject variance but makes the remaining variance more stable and trait-like. Empirically, the strikingly similar patterns of evoked activity during movie watching do not seem to come at a cost to individual identifiability; rather, the most important individual features are preserved or even enhanced during these conditions.

However, the ultimate goal of most individual differences research is not simply to identify a given individual across repeat scans, but to relate variability in brain functional organization to real-world (out-of-scanner) behavior. To this end, the more relevant question is whether connectivity patterns during naturalistic paradigms improve sensitivity to behavioral differences. Previous work using connectivity during traditional tasks to predict fluid intelligence showed that, while certain tasks enjoyed larger advantages than others, all tasks outperformed rest, even when the task state was seemingly unrelated to the construct to be predicted (e.g., even functional connectivity during a finger-tapping task offers improved predictions of fluid intelligence) (Greene et al., 2018). Extrapolating from rest to traditional tasks to naturalistic paradigms, one might predict that naturalistic paradigms would also increase sensitivity to behavior. However, to our knowledge, this has not been directly tested. Furthermore, there has been relatively less work on how brain state affects connectivity-based predictions of traits outside the cognitive domain—for example, emotion and affect. Naturalistic tasks also raise the intriguing possibility of tailoring stimulus content to the construct to be predicted—for example, using a stimulus that evokes feelings of suspense or unease to predict trait or state anxiety, or one that evokes feelings of suspicion to predict paranoia (Finn et al., 2018).

Here, we test the hypothesis that data collected during movie watching improves functional connectivity-based behavior prediction compared to data collected at rest. Using the Human Connectome Project 7T dataset, we demonstrate that trait scores derived from both cognitive and emotion domains are more accurately predicted from movie-watching data, and that this advantage is unlikely to be driven by differences in low-level arousal or data quality between the two conditions. We show that in some cases, both cognition and emotion scores can be predicted from as little as two to three minutes of movie-watching data. Following this, we explore which clip features are associated with prediction accuracy, and find that clips high in social content, as well as those that evoke more variability in gaze position across subjects, tend to yield better predictions. We discuss how these results add to our basic understanding of trait-state interactions in functional connectivity, as well as their practical implications for large-scale efforts in brain-behavior predictive modeling.

## Results

We used fMRI and behavioral data from 176 healthy subjects made available as part of the Human Connectome Project 7T dataset (Van Essen et al., 2013). Functional runs of interest included two resting-state runs and four movie-watching runs, each approximately 15 minutes in duration. While data were acquired in four separate sessions, we focused on the two sessions that contained movie-watching runs (first and fourth). For each subject, REST1, MOVIE1 and MOVIE2 were acquired together in that order in a single session (referred to here as “Session 1”), and REST4, MOVIE3 and MOVIE4 were acquired in that order in a separate session (“Session 2”) on the following day or two days later. During rest runs, subjects were instructed to keep their eyes open and maintain fixation on a central cross. During movie runs, subjects passively viewed a series of four or five video clips, each 1-4 minutes long. Video clips came from both independent and Hollywood films, and varied in their low-level (i.e., audiovisual features) and high-level properties (i.e., semantic content). For a brief description of each clip, see Table 1. For further details on imaging data acquisition, see Materials and Methods.

**Table 1.** Description of individual video clips that comprised each of the four MOVIE runs. MOVIE1 and MOVIE3 contain clips from independent films freely available under a Creative Commons license; MOVIE2 and MOVIE4 contain clips from Hollywood films. * *indicates clip was not included in single-clip CPM analyses due to short length*.

| Run | Clip short name | Full-length film name (year) | Duration (TRs) | Duration (min:sec) | Description |
| --- | --- | --- | --- | --- | --- |
| MOVIE1 | two men | Two Men (2009) | 245 | 04:05 | A man sees another man run past him on a road in rural Australia and muses about his potential motives. Accented English dialogue plus subtitles. |
| MOVIE1 | bridgeville | Welcome to Bridgeville (2011) | 221 | 03:41 | People describe why they love living in small-town America; a collection of scenes from community life. |
| MOVIE1 | pockets | Pockets (2008) | 188 | 03:08 | Close-ups of people holding things they keep in their pockets and describing the items' significance. |
| MOVIE1 | overcome* | Inside the Human Body (2011) | 63 | 01:03 | Inspirational montage of people who have overcome physical disabilities. |
| MOVIE1 | testretest1* | 23 Degrees South (2011); LXIV (2011) | 83 | 01:23 | Concatenation of brief (1-3 s) clips depicting a variety of people, objects, and scenes. |
| MOVIE2 | inception | Inception (2010) | 228 | 03:48 | Two characters explore a dream world and learn its surprising physical and emotional characteristics. |
| MOVIE2 | social net | The Social Network (2010) | 259 | 04:19 | Mark Zuckerberg's disciplinary hearing at Harvard and its aftermath (fictional portrayal). |
| MOVIE2 | ocean's 11 | Ocean's Eleven (2001) | 250 | 04:10 | Danny Ocean and his accomplices meet to plot their Vegas casino heist. |
| MOVIE2 | testretest2* |  | 83 | 01:23 | (same as testretest1) |
| MOVIE3 | flower | Off The Shelf (2008) | 180 | 03:00 | A flower escapes its pot and goes on a journey through the neighborhood. |
| MOVIE3 | hotel | 1212 (year unavailable) | 185 | 03:05 | A man and woman have a metaphysical encounter in a hotel room. |
| MOVIE3 | garden | Mrs. Meyer's Clean Day (2013) | 203 | 03:23 | Documentary about an urban vegetable garden serving community needs. |
| MOVIE3 | dreary | Northwest Passage (year unavailable) | 143 | 02:23 | Montage of dreary landscapes and abandoned structures set to eerie music. |
| MOVIE3 | testretest3* |  | 83 | 01:23 | (same as testretest1) |
| MOVIE4 | home alone | Home Alone (1990) | 234 | 03:54 | Main character walks through the house, realizing he is on his own. |
| MOVIE4 | brokovich | Erin Brockovich (2000) | 231 | 03:51 | Erin Brokovich meets with a plaintiff and then visits a legal office with her young children in tow. |
| MOVIE4 | star wars | The Empire Strikes Back (1980) | 255 | 04:15 | Scene from Rebel base on an icy planet; Leia and Han Solo argue. |
| MOVIE4 | testretest4* |  | 83 | 01:23 | (same as testretest1) |

The HCP makes available extensive phenotyping data for each subject. We were interested in how well functional connectivity during both rest and movies could predict trait behaviors in two broad domains: cognition and emotion. From the individual measures in each domain—many of which are highly correlated with one another—we derived a single score for each subject using the top component from a principal components analysis (one per domain). To avoid dependence between the training and test sets, principal components were learned using an independent set of subjects (n = 1,022; those scanned only at 3T) and this transformation was applied once to all 7T subjects. See Fig. 1 for loadings of individual measures onto these components, and Tables S1 and S2 for full names and constructs for each measure. Briefly, higher scores on the first principal component in the cognitive domain (henceforth referred to as “cognition score”) were associated with better performance on tasks measuring reading ability, vocabulary, and fluid intelligence (Fig. 1a); higher scores on the first principal component in the emotion domain (“emotion score”) were associated with higher self-reported life satisfaction, emotional support, and positive affect, and lower sadness and perceived stress (Fig. 1b). Cognition and emotion scores were not strongly correlated across subjects (r_174_ = 0.11, p = 0.13; Fig. 1c), suggesting that the two domains are largely independent. These two scores served as targets for connectivity-based prediction in the analyses that follow. At each cross-validation fold, prior to model training and testing, scores were residualized with respect to potentially confounding variables (head motion and time of day; see Methods and Supplemental Figs. 1 and 2).

**Figure 1.**
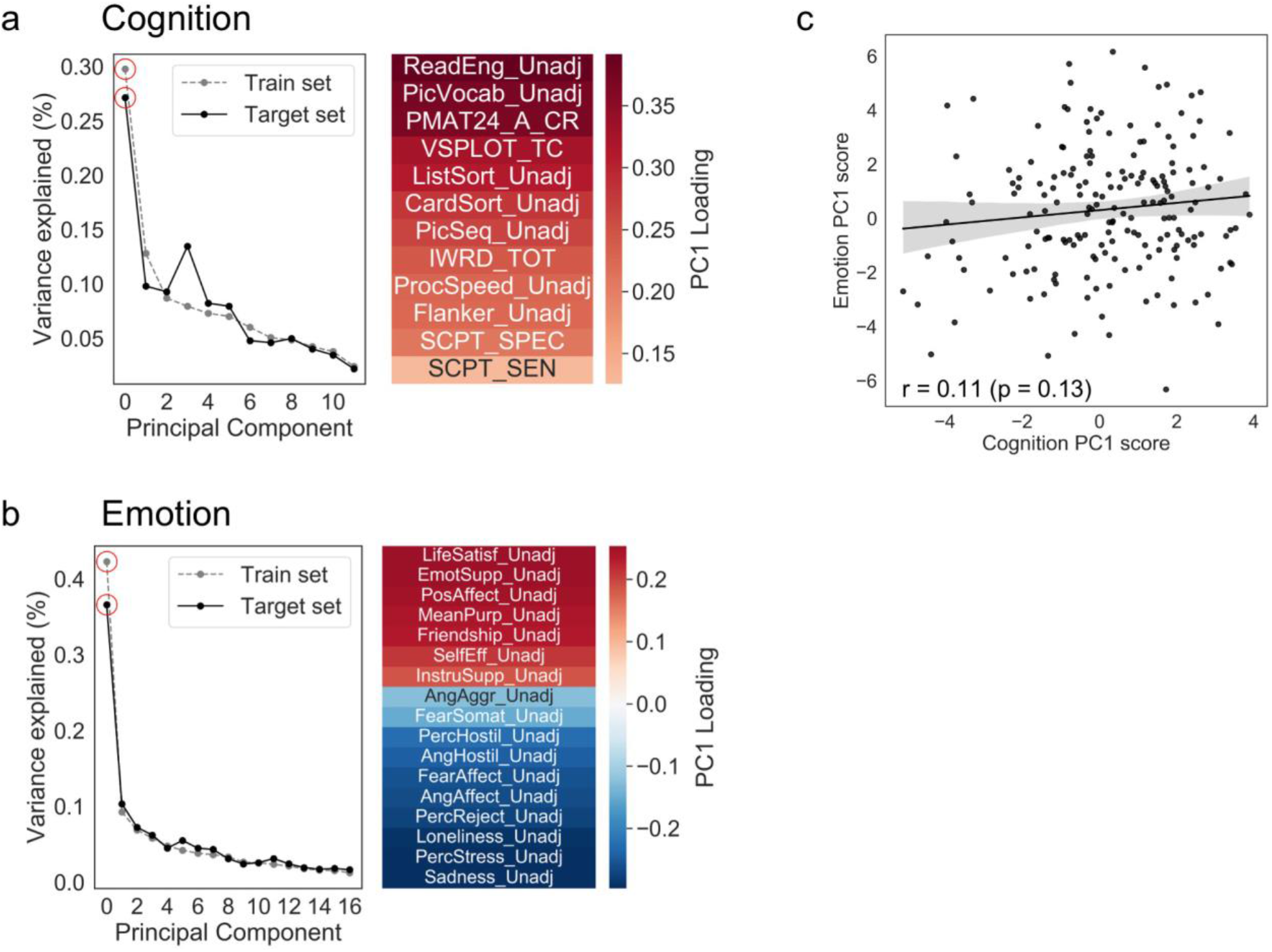
Deriving cognition and emotion scores. Principal components analysis (PCA) was performed on behavioral data from n = 1,022 training subjects (those not scanned at 7T) using measures in the a) cognition and b) emotion domains, and then this learned transformation was applied to the target set of n = 176 subjects in the 7T dataset to derive a cognition and emotion score for these subjects. In the cognition domain, the first principal component explained 30% and 27% of the variance in the training and target sets, respectively, and in the emotion domain, the first principal component explained 42% and 37% of the variance in the training and target sets (red circles on scree plots). The heatmaps show the loadings of individual measures onto the first component in each domain. c) Cognition and emotion scores were not significantly correlated across subjects in the target set, indicating that these two constructs are separable. See Tables S1 and S2 for full variable names and measured constructs for the variables entered into each PCA.

### Movie-watching outperforms rest for prediction of behavior traits

Our main goal was to compare how well cognition and emotion scores could be predicted based on functional brain connectivity during rest or movie watching. REST runs were 900 TRs, or 15:00, in duration, while MOVIE runs ranged from 769-800 TRs (12:49-13:20) following removal of rest periods between clips (see Methods). For each subject for each run, we created a whole-brain functional connectivity matrix by calculating the Pearson correlation of activity timecourses between each pair of nodes in a predefined 268-node atlas (Shen et al., 2013). We used connectome-based predictive modeling (CPM; Finn et al., 2015; Shen et al., 2017) to build and test models trained to predict an individual’s behavioral score from their functional connectivity matrix. Briefly, CPM is a fully cross-validated approach in which a linear model is built to relate connectivity strength in selected features (i.e., connections, or “edges”) to behavioral score within a training set, and then this model is applied to data from subjects in a test set to generate a predicted behavioral score. Here, we used 100 iterations of 10-fold cross-validation. Model accuracy was assessed using Spearman (rank) correlation between predicted and observed scores across all subjects, resulting in one accuracy (*r_s_* value) for each of the 100 iterations.

Cognition scores could be predicted with significant accuracy from all four MOVIE runs (Fig. 2a): MOVIE1 (median r = 0.29, permutation-based p = 0.0009), MOVIE2 (r = 0.41, p = 0.0001), MOVIE3 (r = 0.23, p = 0.011), and MOVIE4 (r = 0.17, p = 0.048), as well as from REST1 (median r = 0.26, p = 0.003), but not from REST4 (r = 0.12, p = 0.13). Within the first session, prediction accuracy was higher for both MOVIE runs than for the REST runs (MOVIE1 versus REST1: Mann-Whitney U-test, effect size [f] = 0.79, p < 10^-12^; MOVIE2 versus REST1, f = 1.0, p < 10^-34^), and was higher for MOVIE2 than for MOVIE1 (f = 1.0, p < 10^-34^). Similarly, within the second session, prediction accuracy was higher for both MOVIE runs than for the REST run (MOVIE3 versus REST4: f = 0.97, p < 10^-31^; MOVIE4 versus REST4: f = 0.84, p < 10^-17^). Also within the second session, accuracy was higher for MOVIE3 than for MOVIE4 (f = 0.84, p < 10^-16^).

**Figure 2.**
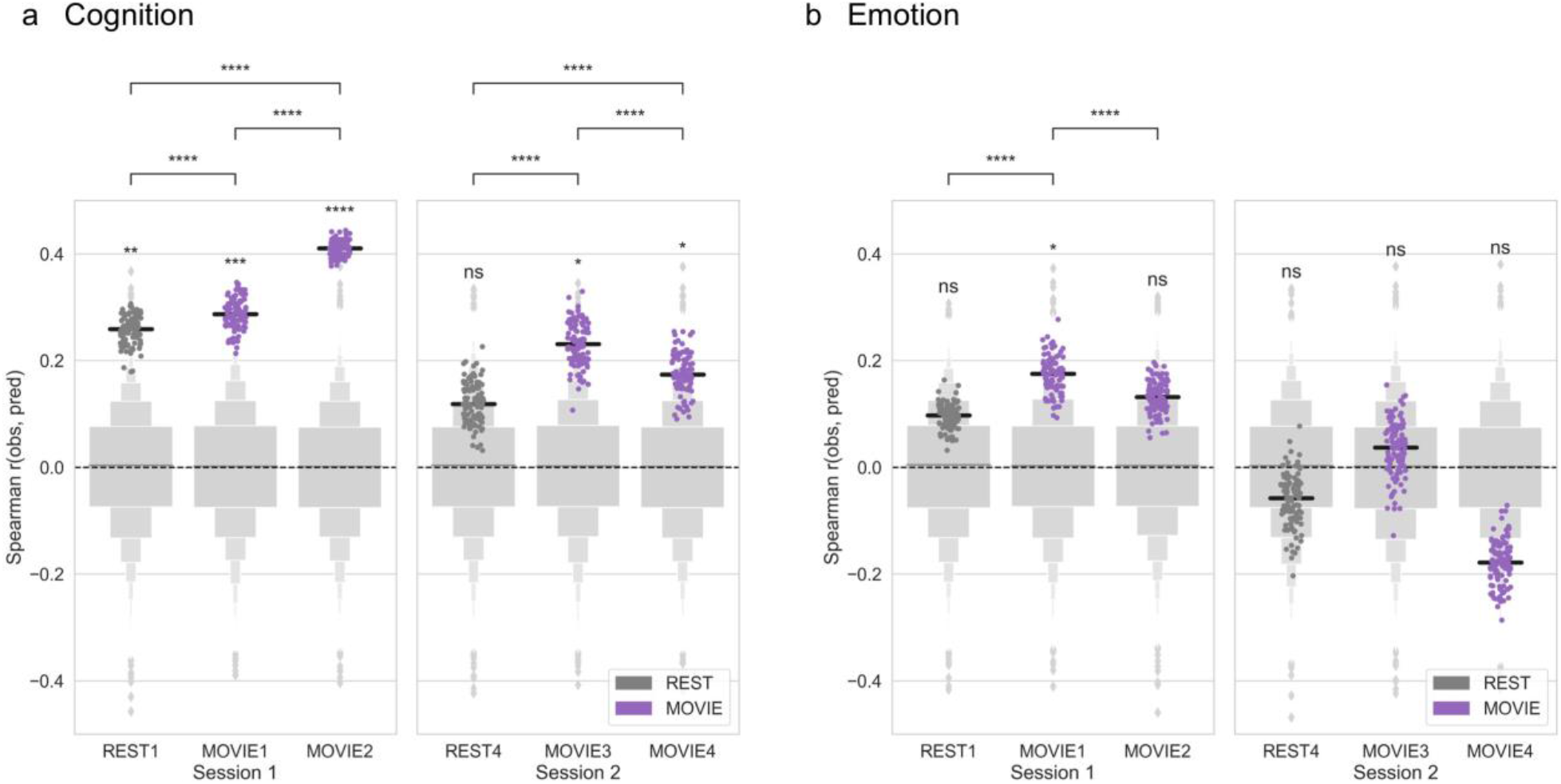
Movie-watching outperforms resting state for functional connectivity-based prediction of behavior. Connectome-based predictive models were trained to predict a) cognition or b) emotion score from resting-state (dark gray) or movie-based (purple) functional connectivity. Accuracy was measured as the Spearman [rank] correlation between predicted and observed scores (*y* axis). Dots show results from 100 iterations of 10-fold cross-validation (true models). Light gray boxen plots show null distribution from 10,000 permutations in which behavior scores and connectivity matrices were randomized across subjects. Black horizontal line denotes median accuracy for true models. Statistical significance for each run was calculated by comparing the median of the true models to the null distribution. Differences in accuracy between runs were assessed using Mann-Whitney U tests. ns: p > 0.05; *p < 0.05; **p < 0.01; ***p < 0.001; ****p < 0.0001.

Emotion scores, on the other hand, could only be predicted from MOVIE1 (r = 0.18, p = 0.049; Fig. 2b). Predictions from REST1 and MOVIE2 did not reach significance, and none of the runs in the second session gave significant predictions (all p > 0.1). Prediction accuracy was higher for MOVIE1 than for REST1 (Mann-Whitney U-test, f = 0.97, p < 10_-30_). However, certain individual clips in both MOVIE1 and MOVIE2 gave significant predictions of emotion scores, as described in the section *Predictions based on individual clips* below.

### Advantage for movies is not due to low-level arousal

Can the advantage for movies be explained by differences in arousal between the two states? In other words, are people simply more likely to be drowsy or asleep during rest, leading to a drop in prediction accuracy? To answer this question, we used eye-tracking data to compare three metrics of alertness across the runs of interest: percentage of TRs with valid eye-tracking data (as a proxy for eyes-open time), blink rate, and median blink duration.

Within session 1, the percentage of TRs with valid-eye tracking data was generally high (approximately 82 percent across all runs), and there were no significant differences in this metric between REST1 and MOVIE1 (paired t-test: t_139_ = −1.48, p = 0.14) or REST1 and MOVIE2 (t_138_ = −0.55, p = 0.59; Fig. 3a). Within session 2, percentages were also high (approximately 82-86 percent). The percentage of TRs with valid data was slightly lower in REST4 than in MOVIE3 (t_143_ = −2.17, p = 0.03), but there was no significant difference between REST4 and MOVIE4 (t_144_ = −1.16, p = 0.25; Fig. 3a).

**Figure 3.**
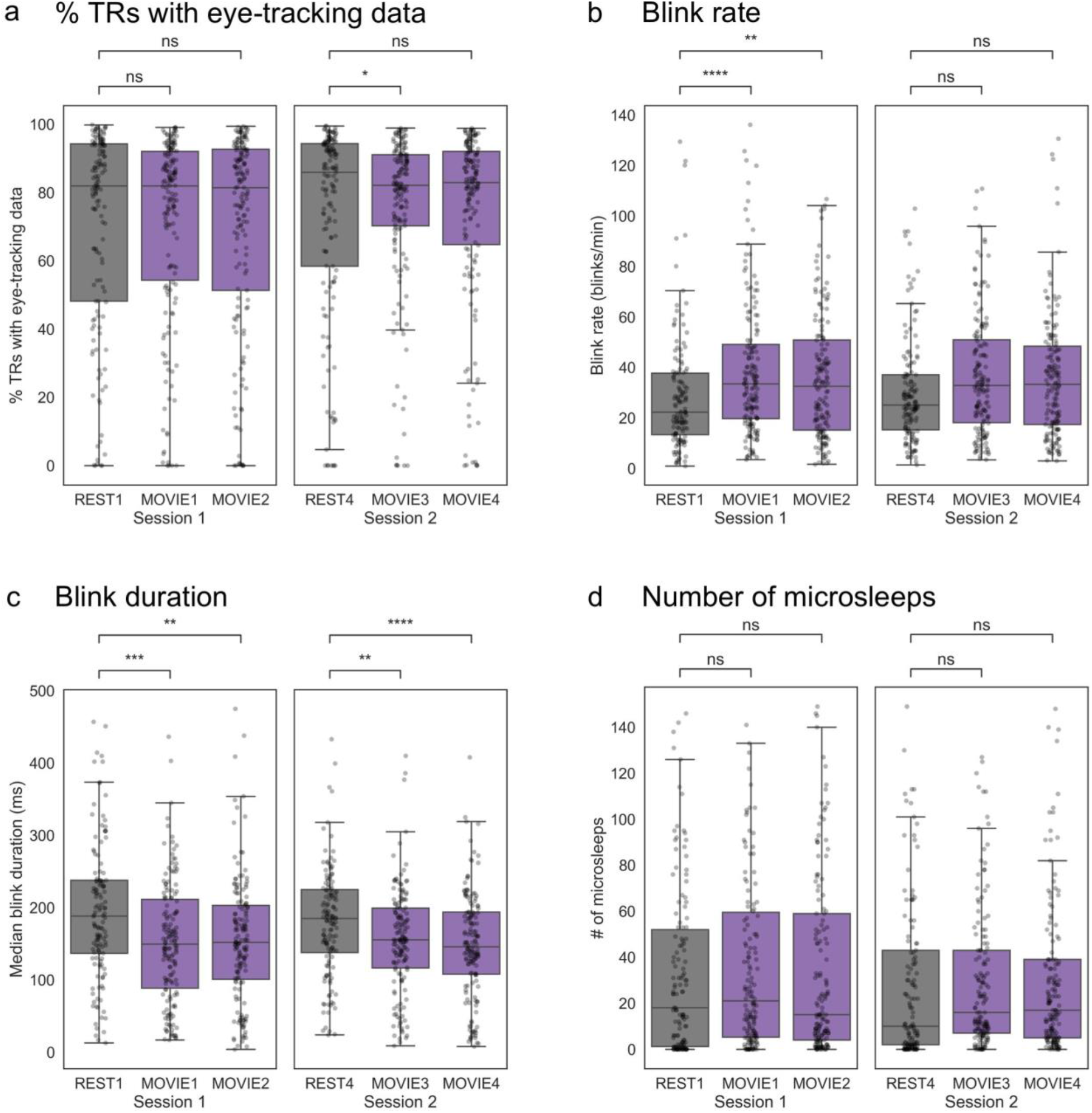
Differences in eye-tracking-based measures between rest and movie runs. Per-run distributions of four metrics are shown: a) percentage of TRs with valid eye-tracking data (a proxy for eyes-open time), b) blink rate, c) blink duration, and d) number of microsleeps (defined as blinks > 1 s in duration). In the box plots, the center line represents the median, while the box extent represents the interquartile range (IQR; 25^th^ – 75^th^ percentile). The whiskers extend to 1.5*IQR. Individual-subject data are overlaid as semi-transparent dots. Note that data are inherently paired (with subject as the repeated measure), but are not displayed as such for reasons of visual clarity. However, all between-run comparisons were conducted using paired t-tests on subjects with data available for both scans in a given pair. n.s., not significant (p > 0.05); *p < 0.05; **p < 0.01; ***p < 0.001; ****p < 0.0001.

We next examined data on blinks. Subjects tended to blink less during REST runs than during MOVIE runs in session 1 (REST1 versus MOVIE1: t_133_ = −5.68, p < 0.0001; versus MOVIE2: t_132_ = −3.14, p = 0.002; Fig. 3b) and, to a lesser extent, session 2 (REST4 versus MOVIE3: t_136_ = −1.95, p = 0.05; versus MOVIE4: t_136_ = −1.76, p = 0.08; Fig. 3b), and blinks were longer in duration during REST runs (REST1 versus MOVIE1: t_133_ = 4.0, p < 0.0001; versus MOVIE2: t_132_ = 2.79, p = 0.006; REST4 versus MOVIE3: t_136_ = 3.1, p = 0.003; versus MOVIE4, t_136_ = 4.4, p < 0.0001; Fig. 3c). Increased blinking during movie-watching may be related to increased saccades, since blinks often accompany saccadic gaze shifts (Evinger et al., 1991). (Participants were instructed to fixate on a central crosshair during rest, while they were free to move their gaze naturally during movie watching.) However, blinks during rest runs were not so long as to imply that subjects were sleeping. “Microsleeps” are typically defined as an eye closure lasting longer than 1000 ms (1 s). In general, microsleeps were rare—the median number across subjects ranged from 10 (REST4) to 21 (MOVIE1)—and there were no differences in number of microsleeps between conditions (all p > 0.23; Fig. 3d). See Fig. S4 for correlations between blink-related measures.

Despite the differences between rest and movie-watching, blink rate and duration were consistent with typical wakeful ranges during both states. Median blink rate across subjects ranged from 22.3 (REST1) to 33.4 (MOVIE1) per minute, which falls within previously published values for spontaneous blink rates (e.g., Bentivoglio et al., (1997), 6-40 per minute; Karson et al. (1983): 23 ± 15 per minute). Median blink duration ranged from 155.5 ms (MOVIE1) to 188.0 ms (REST1), again falling within the typical reported range for the alert state, which is 100-400 ms. Thus, despite some differences in blink patterns, there was no strong evidence that major differences in arousal across conditions are responsible for the observed differences in prediction accuracy.

### Anatomy of predictive networks differs across states

We next examined which functional connections were most important to the predictive models, and whether these differed across states. (Because only MOVIE1 gave significant predictions of emotion score, we restricted these analyses to cognition score to permit comparisons across runs.) Restricting our analysis to connections, or edges, that were selected in at least 90 percent of model iterations, we found significant overlap between nearly all pairs of runs (Fig. 4a), but overlap was higher for runs of the same state than across states (mean overlap within- and across-state, respectively: 36 and 14 edges, p value for difference = 0.0007 according to a Mann-Whitney U test; visible as two distinct clusters in Fig. 4a). This suggests that while some edges are always associated with behavior regardless of state, there is more consistency in the edges associated with behavior within a given state, even across runs and scan sessions.

**Figure 4.**
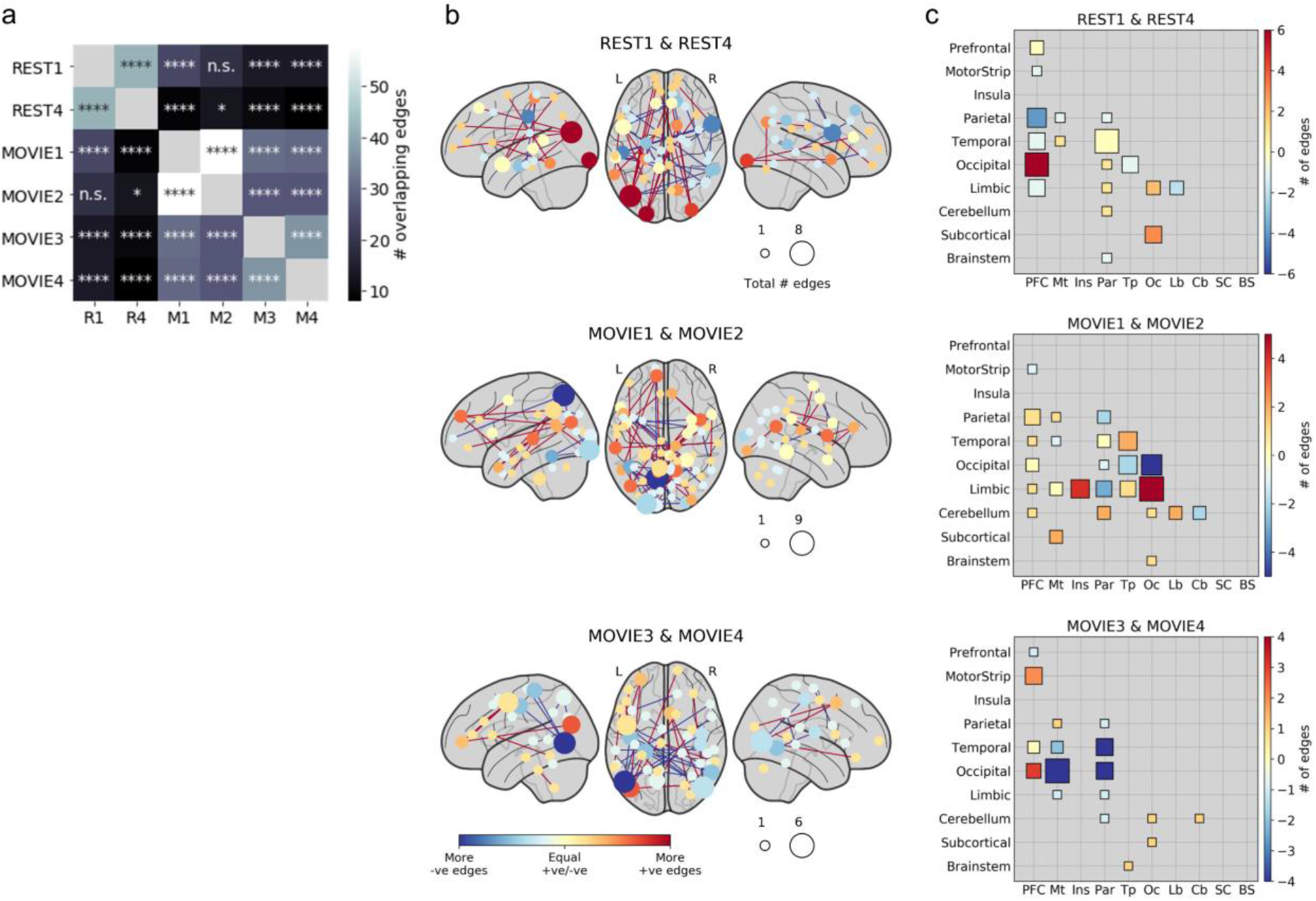
Anatomy of predictive networks across states. a) Overlap between most consistent edges (i.e., those selected in at least 90% of all model iterations) across runs. Significance of overlap was assessed with the hypergeometric cumulative distribution function. n.s., not significant (p > 0.05); *p < 0.05; **p < 0.01; ***p < 0.001; ****p < 0.0001 (Bonferroni corrected). Overlap was higher within states than across states (Mann-Whitney U test, p = 0.0007). b) Nodewise visualization of most consistently selected edges across pairs of runs of the same state (REST or MOVIE). Nodes are sized according to the sum of edges in the positive and negative networks (such that larger nodes had more edges overall), and colored according to the difference between edges in the positive and negative networks (such that red nodes had mostly positive edges and blue nodes had mostly negative edges). “Positive” refers to edges positively correlated with behavior, while “negative” refers to edges inversely correlated with behavior. c) Lobewise visualization of most consistently selected edges across pairs of runs. Size and color scheme are similar to (b). Diagonal depicts within-lobe connections. PFC, prefrontal cortex; Mt, motor strip; Ins, insula; Par, parietal; Tp, temporal; Oc, occipital; Lb, limbic; SC, subcortical; BS, brainstem.

Next, we visualized the most consistently selected edges in two ways: first, by displaying individual nodes in brain space according to their numbers of positive and negative edges (Fig. 4b), and second, by summarizing positive and negative networks by macroscale brain region (i.e., lobe; Fig. 4c). (Note that here, “positive” and “negative” refer to the sign of an edge’s association with behavior, not to its raw correlation strength.) As is typical of a data-driven approach, predictive networks were widely distributed across the brain, with no single dominant anatomical pattern. However, some trends emerged. Stronger connectivity between prefrontal and occipital cortex predicted higher cognition score in both rest runs, and to some extent in the session 2 movie runs (i.e., MOVIE3 and MOVIE4), but were less important in the session 1 movie runs (i.e., MOVIE1 and MOVIE2). Temporal-occipital connectivity was strongly negatively associated with cognition score in MOVIE2, but played a much lesser role in predictive networks in the other runs.

The individual nodes with the most connections in predictive networks also differed across states and sessions. At rest, a node in left visual association cortex (Brodmann’s area 19) had the highest total edges, most of them involving prefrontal cortex and positively associated with cognition score. In the session 1 movie runs, nodes in left parietal cortex (BA 7) and left visual association cortex (BA 18) had the highest total edges, most of them negatively related to cognition score. In the session 2 movie runs, two nearly homologous nodes in left and right visual association cortex had the highest total edges, many of them negatively related to cognition score. Overall, nodes in visual association regions seemed to play an outsize role in predictive networks, though their connection partners and the sign of their association with behavior changed depending on brain state and session. Of note, for the movie runs, it is difficult to disentangle cross-session variability (noise) from effects of stimulus content on selected edges, since each run contained a different set of clips. In the following section, we perform prediction based on single video clips to explore how stimulus content affects prediction accuracy and anatomy of predictive networks.

### Predictions based on individual clips

To explore if and how prediction success varies across stimuli, we tested how well behavioral scores could be predicted from functional connectivity during single video clips. Clips varied in duration from 1:03 to 4:19 min:sec, and previous work has shown that longer acquisitions improve estimation of individual signal (Laumann et al., 2015; Noble et al., 2017). For an unbiased comparison across clips, we truncated all clip timecourses to match the duration of one the shortest clips (143 TRs, or 2:23) and constructed clip-specific functional connectivity matrices for all subjects. We then performed CPM using data from each clip individually to predict cognition or emotion score.

Despite the limited amount of data, several clips yielded significant predictions of behavior (Fig. 5). Cognition scores could be predicted using data from 9 out of 13 clips (“two men”: median r = 0.18, [p = 0.047]; “pockets”: 0.22 [0.017]; “inception”: 0.22 [0.02]; “social net”: 0.38 [0.0001]; “ocean’s 11”: 0.34 [0.0003]; “garden”: 0.18 [0.04]; “home alone”: 0.20 [0.03]; “brokovich”: 0.19 [0.04]; “star wars”: 0.21 [0.02]; Fig. 5a). Emotion scores could be predicted using data from 3 out of 13 clips (“pockets”: median r = 0.22, p = 0.013; “social net”: 0.24 [0.01], and “oceans”: 0.22 [0.01]; Fig. 5b). Similar to the full runs, prediction was overall better for cognition scores than emotion scores; as expected given the whole-run results, no individual clips in MOVIE3 or MOVIE4 yielded significant predictions of emotion score.

**Figure 5.**
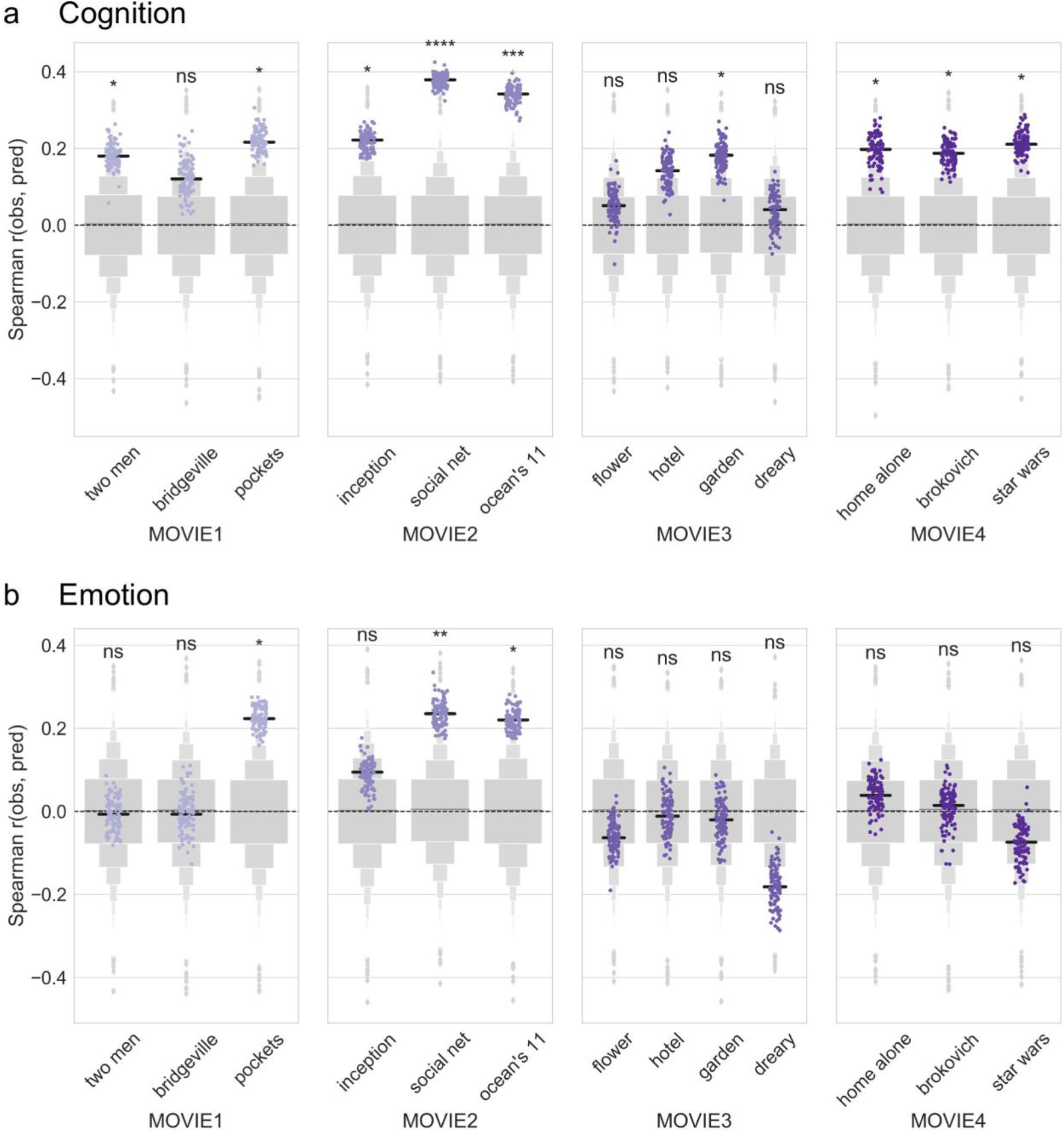
Predictions based on individual video clips. Connectome-based predictive models were trained to predict a) cognition or b) emotion score from individual video clips (*x* axis) in each of the four movie runs. Accuracy was measured as the Spearman [rank] correlation between predicted and observed scores (*y* axis). Dots show results from 100 iterations of 10-fold cross-validation (true models). Light gray boxen plots show null distribution from 10,000 permutations in which behavior scores and connectivity matrices were randomized across subjects. Black horizontal line denotes median accuracy for true models. Statistical significance was calculated by comparing the median of the true models to the null distribution. ns: p > 0.05; *p < 0.05; **p < 0.01; ***p < 0.001; ****p < 0.0001.

We also assessed prediction success for individual clips by comparing accuracies from each clip to a matched block of resting-state data from the REST run in the same session. (For example, the clip “pockets” began at 8:46 into the MOVIE1 run, so its corresponding matched rest block was taken from data beginning at 8:46 into the REST1 run. As above, data was matched for duration at 2:23 min:sec, or 143 TRs.) Overall, 10 and 9 clips (out of 13) outperformed their matched rest block for prediction of cognition and emotion score, respectively (Fig. S3).

We next asked whether high-performing clips were consistent across domains. In other words, if a clip performs well for predicting cognitive score, does it also perform well for predicting emotion score? We correlated median prediction accuracy for cognitive and emotion scores across clips, restricting our analysis to clips in the first session (MOVIE1 and MOVIE2, since no clips in the second session yielded accurate predictions of emotion score). Prediction accuracies were correlated at r_s_ = 0.83 (p = 0.04), indicating that the same clips were most successful for both the cognitive and emotion domains (Fig. 6a).

**Figure 6.**
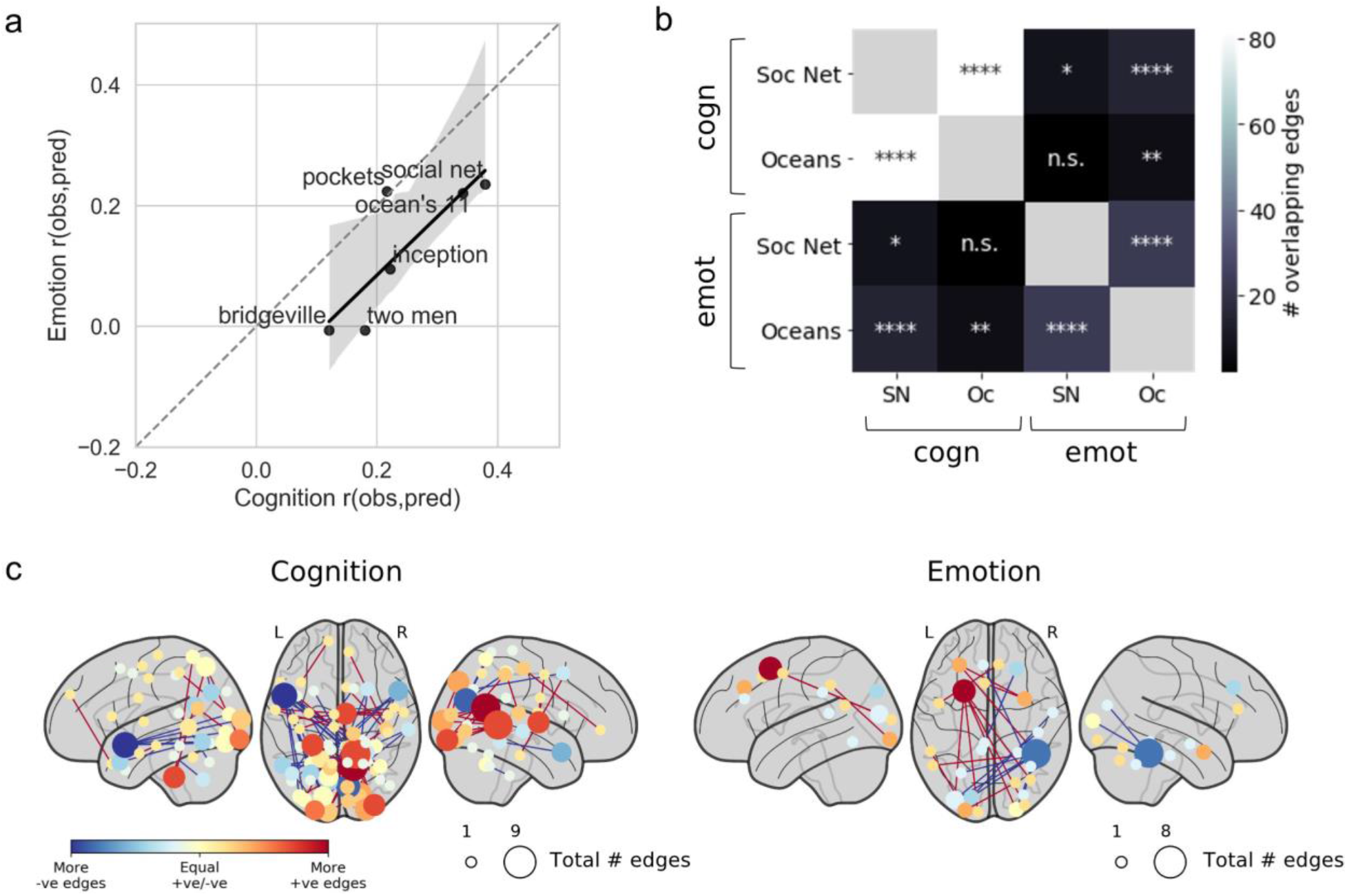
Stimulus- and domain-specificity of predictions. a) Correlation between a clip’s median prediction accuracy for cognitive score (*x* axis) and emotion score (*y* axis), limited to clips acquired in session 1 (since no clips in session 2 gave significant predictions of emotion score). Spearman r = 0.83 (p = 0.04). b) Overlap between consistently selected edges across the two most successful clips (inner label) and domains (outer label). SN, social net; Oc, ocean’s 11. c) Nodewise visualization of consistently selected edges across same two clips in each domain (cognition and emotion). Nodes are sized and colored as in Fig. 4. ns: p > 0.05; *p < 0.05; **p < 0.01; ***p < 0.001; ****p < 0.0001.

For the most successful clips, are the same edges selected for both the cognitive and emotion predictive models, or do the informative edges differ by domain? To answer this, we assessed the overlap between the consistently selected edges in the cognition and emotion models in the two clips that performed best for both domains, Ocean’s 11 and Social Network. Models trained to predict the same behavior from different clips had more overlapping edges than models trained on the same clip to predict different behaviors (Fig. 6b). In other words, similar networks were selected to predict cognitive score from both Ocean’s 11 and Social Network (overlap depicted in Fig. 6c, left panel), but different networks were selected to predict emotion score (overlap depicted in Fig. 6c, right panel). Therefore, the same clip may perform well across domains, but different edges may be important to the model in each particular domain.

### Stimulus features related to prediction accuracy

Which clip features are associated with more accurate predictions? Understanding how clip content relates to prediction accuracy can shed light on why certain clips are more successful than others, which may help guide stimulus selection for future studies. (For these analyses, we again restricted our analyses to predictions of cognitive score because too few clips gave significant predictions of emotion score to draw conclusions.)

We first used semantic category labels available for each clip that were created by hand using the WordNet semantic taxonomy (Huth et al., 2012) and made available as part of the HCP dataset. These annotations contains 859 distinct object (noun) and action (verb) categories. Using partial least-squares regression, we identified one component of these labels that explained 96 percent of the variance in median prediction accuracy across clips (Fig. 7a). Categories with strong positive loadings on this component (associated with better prediction accuracy) included verbs such as “act” and “talk”, as well as nouns such as “person” and “causal agent”. Categories with strong negative loadings (associated with worse prediction accuracy) were largely nouns rather than verbs, and included objects associated with scenes or landscapes (e.g., “mammal”, “telephone pole”, “building material”, “vegetation”). This suggests that clips with more humans, verbal interaction, and social information yield better behavioral prediction than clips containing mostly nature scenes and less social content.

**Figure 7.**
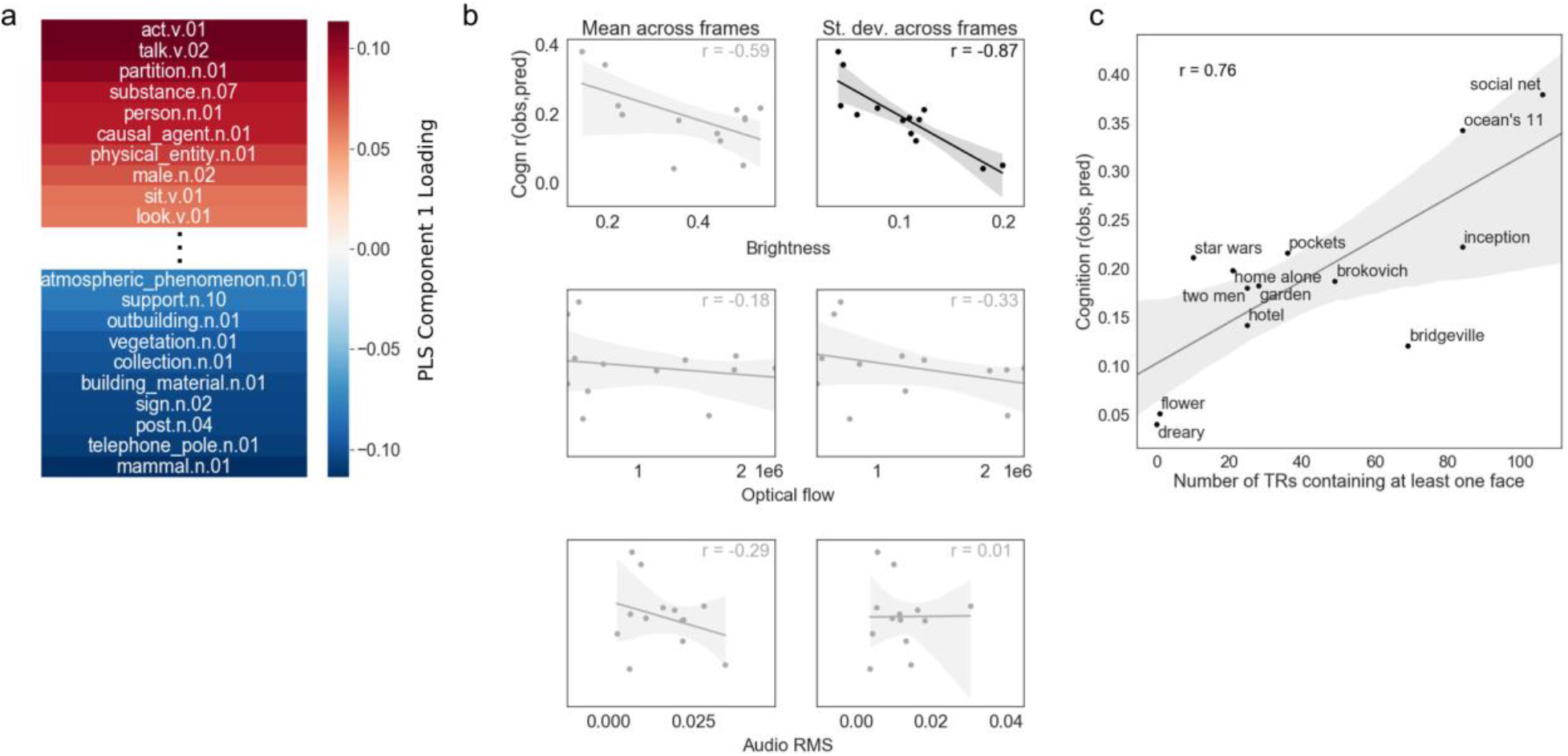
Stimulus features related to prediction accuracy. a) First component from a partial least squares regression relating semantic content to prediction accuracy for cognition score across video clips. Positive weights (red) are associated with better accuracy, while lower weights (blue) are associated with worse accuracy. b) Correlations between prediction accuracy for cognition score (*y* axis) and low-level audiovisual features (*x* axis; from top to bottom row: brightness [luminance], optical flow [between-frame motion], and audio RMS [volume]). Left column is mean across video frames, right column is standard deviation across video frames. Following correction, the only significant relationship was with brightness standard deviation, such that less variance in brightness across a clip was associated with better accuracy (r = −0.87, corrected p = 0.0008). c) Correlation between number of TRs containing at least one face onscreen (a proxy for social content) and prediction accuracy for cognition score (y axis): (r = 0.76, p = 0.003).

In another analysis, we used automated feature extraction (McNamara et al., 2017) to create additional annotations of both low- and high-level features for each clip. Low-level features included brightness (luminance), optical flow (amount of frame-to-frame visual motion), and audio power (root-mean square of sound signal amplitude). To test the hypothesis arising from the WordNet semantic analysis that social content is associated with better prediction, we also labeled clips for the presence of human faces. We then summarized these properties for each clip using both mean and standard deviation across TRs (for low-level features) and number of TRs containing at least one face (for the high-level face feature) and correlated these with median prediction accuracy (Fig. 7b, 7c). Of the low-level features, only brightness standard deviation was significantly correlated with prediction accuracy, such more variance in brightness level across clip frames was associated with lower prediction accuracy (r = −0.87, Bonferroni corrected p = 0.0008). As predicted, number of TRs containing at least one face was positively correlated with median prediction accuracy (r = 0.76, p = 0.003). However, note that some of these features were also correlated with one another (Fig. S5); number of faces was strongly negatively related to brightness standard deviation (r = −0.77), making it difficult to attribute unique variance in prediction accuracy to either feature.

### Cross-subject variance in gaze location is associated with better prediction

One possibility is that participant engagement mediates the relationships between stimulus features and prediction success. In other words, if clips with more social content are simply more engaging, these clips may yield more accurate predictions because they evoke richer cognitive states and/or standardize arousal levels across subjects. In the absence of explicit ratings of engagement or behavioral proxies (e.g., debrief questionnaires or comprehension questions), how might we measure engagement? We reasoned that one index of engagement might be synchrony of gaze location, such that more engaging clips would evoke more similar patterns of eye movements across subjects. To test this hypothesis, for each individual clip, we used eye-tracking data to create gaze-position inter-subject correlation (ISC) matrices by correlating horizontal gaze location across time between each pair of subjects. We then correlated the median gaze-ISC value with cognition prediction accuracy across clips.

Contrary to our hypothesis, median gaze ISC was not correlated with prediction accuracy (r_11_ = 0.06, p = 0.85; Fig. 8a). However, standard deviation of gaze ISC was correlated with prediction accuracy (r_11_ = 0.61, p = 0.03; Fig. 8b), meaning that clips that evoked more variable patterns of eye movements across participants were better predictors. Standard deviation of gaze ISC was not strongly correlated with number of TRs with faces onscreen (r_11_ = 0.32, p = 0.28), suggesting that these two factors related to prediction accuracy are at least partially dissociable.

**Figure 8.**
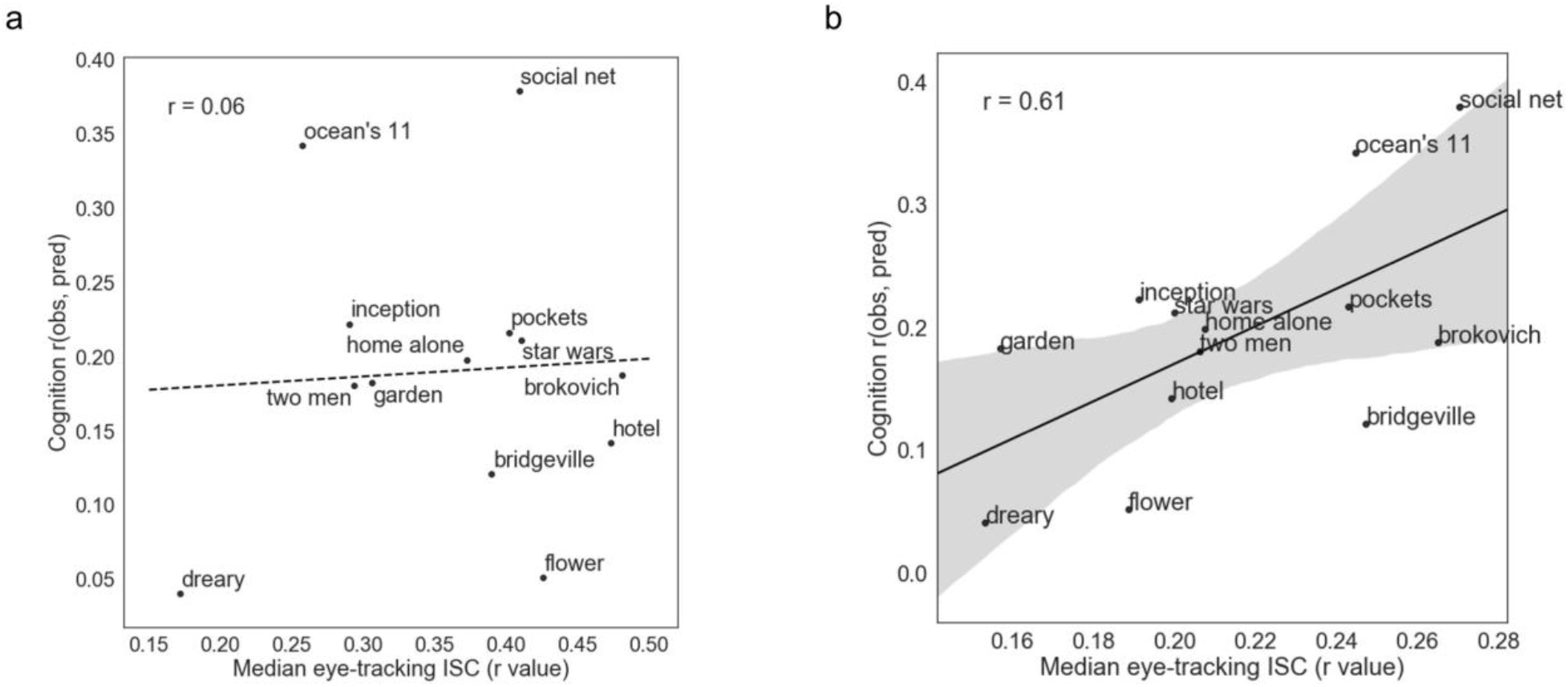
Cross-subject variance in gaze position is associated with better prediction. Relationships across clips between prediction accuracy for cognition score (*y* axis) and measures of inter-subject correlation (ISC) in horizontal gaze position across subject pairs (*x* axis). While there was no significant relationship between median gaze position ISC and accuracy (a), higher accuracy was associated with higher standard deviation of gaze position ISC (b), suggesting that the most successful clips are those that evoke the most variability in gaze trajectories across subjects.

Cross-subject variance in gaze trajectory may lead to higher prediction accuracy if there is a relationship between gaze trajectory, functional connectivity, and trait scores. In other words, subjects who are more similar in their eye movements may also be more similar in their functional connectivity, and if these pairs of subjects are also more similar in their trait scores, this might explain why clips that evoke higher gaze variance are better predictors: they emphasize similarities between specific pairs of subjects, rather than indiscriminately boosting similarity across all pairs. The relationships among gaze trajectory, functional connectivity, and trait-like phenotypes—and whether certain types of content elicit more variance in gaze trajectory across subjects—should be investigated in future work.

## Discussion

Here, we demonstrated that functional connectivity during movie watching outperforms rest for predicting trait-like behavioral scores in both the cognitive and emotional domains. There was no evidence that this effect was driven by differences in overall alertness across states, suggesting that the advantage for movies may stem from a shift in ongoing mental processes rather than low-level arousal changes. Within movie runs, several individual video clips gave successful predictions of trait scores based on only ~2.5 minutes of data. The best-performing clips were those with strong social content (i.e., humans, faces, dialogue), and also tended to evoke more variability in gaze trajectory across subjects. These results have implications for our basic understanding of trait-state interactions in relationships between functional connectivity and behavior, and for practical considerations related to future data collection efforts aimed at predictive modeling of behavior. We discuss these in turn below.

In general, prediction was more successful for the cognitive domain than the emotion domain. This result is perhaps not surprising in the context of prior work, as many studies have demonstrated successful prediction of cognitive ability—most typically, fluid intelligence and/or working memory—from functional connectivity using the HCP dataset (Smith et al., 2013; Finn et al., 2015; Hearne et al., 2016; Ferguson et al., 2017; Greene et al., 2018; Li et al., 2019) and other datasets (Hampson et al., 2006; van den Heuvel et al., 2009; Cole et al., 2012; Greene et al., 2018; Li et al., 2019), but there are comparatively fewer reports of functional connectivity predicting emotional or affective traits out-of-sample, and accuracies tend to be lower for these traits (Kong et al., 2019). One major difference is that the measures comprising the cognition score were performance-based, while those comprising the emotion score were self-reported. Self-report measures can suffer from bias, and may be less biologically valid than task performance. Ongoing efforts to develop behavior-based computational phenotypes for emotional and affective traits (Montague et al., 2012; Patzelt et al., 2018) may lead to measures that are more biologically valid and therefore more readily predicted from functional connectivity data. Another possibility that does not appeal to construct validity is that these measures simply do not have robust correlates in static functional brain connectivity, but rather in other properties of brain structure or function. Still, we observed significant predictions of emotion score using data from one movie run (MOVIE1) and three individual clips (interestingly, two of which were from a different run, MOVIE2) but not from any resting run. This underscores the heightened sensitivity of moviewatching connectivity to even hard-to-predict measures, and suggests that stimulus content may be particularly important for successful emotion prediction.

Why does functional connectivity during movies yield better predictions of trait behaviors than functional connectivity during rest, and why might certain movies—i.e., those with strong social content—perform better than others? Differences in model accuracy using different states (i.e., rest versus movie-watching) and stimuli (i.e., different clips) to predict the same target behaviors suggests that there are trait-state interactions at play, such that movies, and certain movies in particular, enhance individual differences in behaviorally relevant connections. Previous work has investigated how movie-watching and other naturalistic paradigms affect functional connectivity, with reports that relative to rest, movie-watching alters how activity propagates across cortical pathways (Gilson et al., 2018), especially within and between regions related to audiovisual processing and attention (Betti et al., 2013; Demirtaş et al., 2019), and pushes the network community structure into an overall less modular and more integrated state (Betzel et al., 2020). However, it is not clear whether and how these modulations are related to behavioral variability across subjects. In other words, is this reorganization more pronounced in individuals at one end of the phenotypic spectrum, hence the increased sensitivity to trait-level differences for movies? Future work should relate movie-induced changes in individual edge strengths (using, for example, the approach taken by Greene et al. (2020)) and/or network-level properties to behavior, to better understand why and how movie watching boosts sensitivity to phenotypic differences.

How might we interpret the success of social videos in particular? The relationship between a video’s social content (proxied by number of faces onscreen) and its success in predicting cognition score is intriguing, given that this score is comprised of tests of fluid intelligence, working memory, and other constructs that are not explicitly social in nature. Broadly speaking, this fits with the social brain hypothesis that the human brain and its myriad cognitive advances chiefly evolved to handle increasingly complex social structures (Dunbar, 1998). One possibility is that social videos are simply more engaging, and therefore more effective at corralling subjects into a similar brain state, reducing noise and enhancing behaviorally relevant signal among any remaining differences. However, if this were the case, we might expect gaze trajectories to also be more similar across subjects in the most successful videos (reflecting shared attention and processing), and this was not what we found. Instead, more successful videos tended to have more variability in gaze trajectories, such that some pairs of subjects were highly synchronized while others were dissimilar. Perhaps the most successful videos are those that engage people in different ways, and social content is the most likely to evoke different reactions and interpretations across subjects (Finn et al., 2018; Chen et al., 2019; Gruskin et al., 2019; Nguyen et al., 2019). Future work using online or post-hoc measures of stimulus engagement and interpretation could further explore this possibility.

The most successful clips were from Inception, The Social Network, and Ocean’s 11, which are all relatively recent and well-known Hollywood films. This could have conferred additional advantages for at least two reasons (not necessarily mutually exclusive with those presented above). First, many participants might have seen these films in their entirety prior to the scan session, meaning they would have been able to place the short clip into a larger context that may have evoked richer mental representations during viewing compared to an unfamiliar film. (Participants were not asked whether they had seen any of the films previously, but future studies should include this information in debriefing so that these effects can be studied and/or controlled for.) Second, the film industry, and Hollywood-style films in particular, generally gets better over time at using cinematographic techniques to capture audience’s attentions and emotions, boosting engagement (Cutting et al., 2011). Both of these are important factors to consider in designing future studies.

### Limitations

Our study has several limitations worth noting. First, we used a single group parcellation to define nodes. If individual subjects’ functional anatomies are misaligned to this parcellation, it could bias measurements of functional connectivity (Bijsterbosch et al., 2018). Individual- and/or state-specific parcellations (Salehi et al., 2020) could capture additional variance in trait scores (Kong et al., 2019) and/or improve accuracy of CPM-based predictions. (However, we note that this parcellation was defined on resting-state data, which if anything should bias results in favor of rest over movie-watching, and thus work against our hypothesis.) Second, we investigated only one prediction algorithm, connectome-based predictive modeling (Shen et al., 2017). While we chose CPM because it provides a good balance of accuracy, computational tractability, and interpretability, and to be consistent with prior work from our group and others, future work should investigate if other algorithms improve prediction accuracies overall and/or for one condition over another. A third limitation concerns preprocessing choices and potential confounding effects. While we chose to work with the HCP-provided FIX-denoised data with the addition of global signal regression (since this step boosts associations between connectivity and behavior (Li et al., 2019)), it is possible that other preprocessing choices might influence overall accuracy and/or the pattern of results. Similarly, while we attempted to control for collinearity between target behaviors and known confounds (head motion, time of day), it is always possible that insidious effects persist and bias predictions (Siegel et al., 2016).

Finally, perhaps the biggest limitations stem from experimental design and data collection for this study, which was not necessarily designed to test these hypotheses. The order of video clips within runs as well as the order of runs themselves were not counterbalanced, raising the possibility that runs and/or clips in session 2 (which was actually the fourth session in the overall protocol) performed worse for prediction simply due to fatigue effects and not differences in clip content. Within a session, the rest run was always collected before movie runs (though we note again that if anything this is likely to work against our hypothesis, as fatigue generally increases and compliance decreases over the course of an imaging session). Video clips were not explicitly selected to cover a broad space of low-level and high-level features, and as such, there is less range in clip content than might be desired. (For example, most clips come from Hollywood or documentary-style films and contain predominantly social content; most of the clips that buck this trend—i.e., those with predominantly nature scenes and/or little or no dialogue—were confined to a single run, MOVIE3.) Future work should be deliberate in selecting stimuli that vary along certain dimensions of interest and counterbalancing order to provide a clearer picture of how stimulus content affects prediction accuracy.

### Implications for future work

In spite of these limitations and outstanding questions, we believe that the results presented here should encourage more widespread adoption of naturalistic paradigms for studies of brain-behavior relationships. Large-scale data collection efforts might consider including a movie-watching condition in addition to, or perhaps even instead of, resting state. Beyond improving subject compliance, naturalistic paradigms seem to enhance meaningful individual variability in functional connectivity, akin to a “stress test” for the brain (Dubois and Adolphs, 2016; Finn et al., 2017). This may hasten development of biomarkers with real-world applications (Eickhoff et al., 2020).

Of the myriad potential ways to analyze data from naturalistic paradigms, we restricted our approach here to functional connectivity because it can be applied equally to both resting-state and movie-watching data, providing a straightforward way to test for differences between conditions. However, analyses that exploit the presence of a time-locked “ground truth” in naturalistic paradigms, such as inter-subject representational similarity analysis (Finn et al., 2020), an extension of the inter-subject correlation family of approaches (Hasson et al., 2004; Nastase et al., 2019), might prove even more sensitive to trait-level variability in both normative (van Baar et al., 2019) and clinical populations (Hasson et al., 2009; Salmi et al., 2013; Byrge et al., 2015; Bolton et al., 2018; Salmi et al., 2020). Because these and other activation-based approaches allow us to interpret spatiotemporal activity patterns in the context of stimulus features, they open the door to models that are not only predictive (*what* are the relationships between brain function and behavior), but also deepen our understanding of *how* and *why* individuals show distinct brain responses to the same information, and how this relates to trait-level phenotypes.

## Acknowledgments

Data were provided by the Human Connectome Project, WU-Minn Consortium (Principal Investigators: David Van Essen and Kamil Ugurbil; 1U54MH091657) funded by the 16 NIH Institutes and Centers that support the NIH Blueprint for Neuroscience Research; and by the McDonnell Center for Systems Neuroscience at Washington University. This work utilized the computational resources of the NIH HPC Biowulf cluster (http://hpc.nih.gov), and was supported by the National Institutes of Health (grants K99MH120257 and R00MH120257 to E.S.F. and ZIAMH002783 to P.A.B.).

## Data and code availability

Original MRI and behavioral data are publicly available via the Human Connectome Project. Data derivatives (e.g., nodewise timeseries) and code used to conduct the analyses and generate the figures in this manuscript are publicly available in the following repository: https://github.com/esfinn/movie_cpm

## Methods

### Data

#### Subjects

All data used here come from the Human Connectome Project (HCP) 7T release. A total of 184 subjects were scanned at 7T; these were a subset of the approximately 1,200 subjects scanned at 3T. We limited our analysis to subjects who had complete data for all six functional runs of interest as well as complete data for the phenotypic variables of interest (described further below in the sections *fMRI data* and *Behavioral data*, respectively), yielding a set of n = 176. Notably, there were many sets of twins (both mono- and dizygotic) and siblings in this dataset, such that these 176 subjects came from only 90 unique families. Relatedness was taken into account during cross-validation using a leave-one-family-out approach, described further in the section on *Connectome-basedPredictive Modeling*. All subjects were generally healthy young adults between 22-36 years old (mean age = 29.4, standard deviation = 3.3).

#### fMRI data

All fMRI data were acquired on a 7 Tesla Siemens Magnetom scanner at the Center for Magnetic Resonance Research at the University of Minnesota. There were four scan sessions total spread across two or three days; we focus here on the first and last session (which we refer to as session 1 and session 2 here), since these contained the movie-watching runs of interest. REST and MOVIE runs were collected using the same gradient-echo echo-planar imaging (EPI) sequence with the following parameters: repetition time (TR) = 1000 ms, echo time (TE) = 22.2 ms, flip angle = 45 deg, field of view (FOV) = 208 x 208 mm, matrix = 130 x 130, spatial resolution = 1.6 mm3, number of slices = 85, multiband factor = 5, image acceleration factor (iPAT) = 2, partial Fourier sampling = 7/8, echo spacing = 0.64 ms, bandwidth = 1924 Hz/Px. The direction of phase encoding alternated between posterior-to-anterior (PA; REST1, MOVIE2, MOVIE3) and anterior-to-posterior (AP; REST4, MOVIE1, MOVIE4).

Each REST run was 900 TRs, or 15:00 minutes, in length. MOVIE runs 1-4 were 921, 918, 915, and 901 TRs, respectively. However, when calculating functional connectivity during MOVIE runs, we discarded TRs corresponding to the 20-second rest blocks in between video clips (described in more detail in *Stimuli* below). To avoid large changes in BOLD signal at the onset of individual clips that could skew correlations between node timecourses, we excluded the first 10 TRs (10 seconds) of each clip when calculating functional connectivity. On the other hand, to account for hemodynamic delay, we included the 5 TRs (5 seconds) after video offset in the calculation. This lead to effective durations of 775, 800, 769, and 783 for the MOVIE runs. If anything, the reduced duration for movie runs compared to rest should disadvantage movies, working against our hypothesis.

While four REST runs were collected, here we use only data from REST1 and REST4, as these were acquired in the same scan sessions as MOVIE1/MOVIE2 and MOVIE3/MOVIE4, respectively. Within a session, REST runs were always acquired first, followed by the movie runs in a fixed order, such that session 1 consisted of REST1, MOVIE1, and MOVIE3, and session 2 consisted of REST4, MOVIE3, and MOVIE4.

All analyses began with the FIX-denoised data in volume space (e.g., *rfMRI_REST1_7T_PA_hp2000_clean.nii.gz; tfMRI_MOVIE1_7T_AP_hp2000_clean.nii.gz*), which includes standard preprocessing (motion correction, distortion correction, high-pass filtering, and nonlinear alignment to MNI template space (Glasser et al., 2013) plus regression of 24 framewise motion estimates (six rigid-body motion parameters and their derivatives and the squares of those) and regression of confound timeseries identified via independent components analysis (Griffanti et al., 2014; Salimi-Khorshidi et al., 2014). In addition to this preprocessing, we calculated the average whole-brain signal at each TR using an HCP-provided per-subject, per-run mask (*brainmask_fs.1.60.nii.gz*) and regressed this from the FIX-denoised images, in light of prior work (Li et al., 2019) and our own unpublished observations that global signal regression strengthens the association between functional connectivity and behavior.

#### Stimuli

During REST runs, subjects were instructed to keep their eyes open and maintain relaxed fixation on a projected bright cross-hair on a dark background.

During MOVIE runs, subjects passively viewed a series of video clips with audiovisual content. Each MOVIE run consisted of 4 or 5 clips, separated by 20 s of rest (indicated by the word “REST” in white text on a black background). Two of the runs, MOVIE1 and MOVIE3, contained clips from independent films (both fiction and documentary) made freely available under Creative Commons license on Vimeo. The other two runs, MOVIE2 and MOVIE4, contained clips from Hollywood films. The last clip was always a montage of brief (1.5 s) videos that was identical across each of the four runs (to facilitate test-retest and/or validation analyses). For brief descriptions of each clip, see Table 1. Audio was delivered via Sensimetric earbuds.

Clips varied in length from 1:03 to 4:19 min:sec. Because having more data typically boosts accuracy in both individual identification and behavioral prediction, for analyses where we wished to compare prediction performance across clips, we truncated data from each clip to the length of one of the shortest clips, which was 143 TRs (2:23 min:sec). The very shortest clip was only 1:03 in duration, and the test-retest clips at the end of each run were 1:23 in duration. Because truncating all clips to 83 TRs would have severely limited the data available to the model, we omitted these shortest clips from the comparisons.

##### Behavioral data

HCP provides a large number of phenotypic measures from a variety of domains. We focused on traits in two domains: 1) cognition, and 2) emotion/affect. To avoid incurring a multiple-comparisons issue by training models for each individual measure, and because many measures within a domain are correlated with one another, we first performed principal components analysis (PCA) to reduce the dimensionality of the data. Importantly, given that the subjects scanned at 7T are a subset of those scanned at 3T, we learned the principal components on the complement of the 7T subset—in other words, the subjects scanned at 3T but not at 7T (n = 1,022)—and applied this PC-based transformation to data from the 7T subjects to derive PC scores for each 7T subject. Thus we were able to calculate these summary scores once at the start of our analysis pipeline without incurring circularity, or leakage of information from the test set into the training set. (Otherwise, we would have had to reperform the PCA step at each fold of each cross-validation procedure, resulting in slightly different transformations each time.)

PCA was performed separately for the cognition and emotion domains. Individual measures were normalized to have zero mean, unit variance in the training set, and this same normalization (using the mean and standard deviation from the training set) was applied to the test set prior to PCA transformation. Variables entered into the cognition PCA included measures from the NIH Cognition Toolbox as well as additional measures from other instruments classified in “cognition” by the HCP. Variables entered into the emotion PCA were scales from the NIH Emotion Toolbox, a self-report battery assessing A full list of measures entered into each PCA is provided in Supplementary Tables 1 and 2.

#### Eye-tracking data

Eye tracking was acquired during both REST and MOVIE runs using an EyeLink S1000 system (SR Research). We extracted eye-tracking data from two files: 1) the HCP-provided session summary for each subject for each run (e.g., *100610_7T_MOV1_eyetrack_summary.csv*), which provides metadata and quality control measures, and 2) the raw EyeLink log files (e.g., *100610_7T_MOV1_eyetrack.asc*), which provide horizontal position, vertical gaze position, pupil size measures for each timepoint, as well as tags corresponding to blink onset and offset. Of the 1,056 runs of interest (176 subjects x 6 runs each), valid eye-tracking data were available for 931 runs. While most of these (n = 835) had a sampling rate of 1000 Hz, a few (n = 96) had a sampling rate of 500 Hz. All 931 sessions were used for analyses of blinks, but analyses of inter-subject correlation in gaze position were limited to runs with sampling rate of 1000 Hz.

For analyses of blinks, while all available data points are shown in the boxplots in Fig. 3, due to the paired nature of the comparisons, the input to statistical tests was limited to subjects that had valid data for both runs in each pair of interest. Between-clip rest blocks were not removed from MOVIE runs in analyzing blinks.

In calculating inter-subject correlations in gaze position for individual video clips (analyses shown in Fig. 8), we discarded the first 5 seconds after video onset, to avoid biases from large jumps in gaze position at the start of a video.

### Mitigating confounds

Head motion produces well-known artifacts in functional connectivity. To determine if and how head motion might confound our analyses, we assessed whether and to what extent 1) head motion differed across runs, and 2) head motion was correlated with target behavior scores, using mean framewise displacement across TRs as our measure of head motion (*Movement_RelativeRMS_mean.txt*).

In session 1, MOVIE1 had higher motion than REST1 (paired t-test: t_175_ = 5.3, p < 10^-6^; note that this is the opposite of what might be the expected direction), but there was no difference between REST1 and MOVIE2 (t_175_ = 1.0, p = 0.32). In session 2, REST4 had higher motion than both MOVIE3 (t_175_ = 8.9, p < 10e^-15^) and MOVIE4 (t_175_ = 3.3, p = 0.001) (Fig. S1a).

Similar to previous reports in this (Siegel et al., 2016) and other datasets, head motion was negatively correlated with cognition score. This was true in all six runs (see Fig. S1b for correlation coefficients and p-values). However, critically, the magnitude of this correlation did not differ across any pair of runs (all p > 0.11, Steiger’s Z test). This makes it unlikely that differences in prediction accuracy across runs are influenced by the relationship between head motion and cognition score. Head motion was not correlated with emotion score in any of the six runs (all p > 0.16; Fig. S1c).

Given reports that time of day can affect measurements of functional connectivity, we also assessed differences across runs in the time of day they were acquired. Acquisition time for session 1 followed a bimodal distribution, with some subjects being scanned in mid-to late morning, while others were scanned fairly late in the evening (centered roughly around 20:00, or 8:00pm). Session 2 followed a unimodal distribution centered around noon, but still with considerable variability across subjects (Fig. S2a). To determine whether time-of-day effects might pose a confound for our behavioral prediction analyses, we correlated time of day with behavior score across subjects. For runs in session 1, time of day was negatively correlated with cognition score, such that subjects who were scanned later in the day tended to have lower scores (Fig. S2b). These correlations were particularly surprising given that to the best of our knowledge, behavioral data acquisition took place at Washington University in St. Louis before participants were flown to Minneapolis for the 7T portion of the study at University of Minnesota. Therefore the behavioral and fMRI data acquisitions were likely separated by a period of at least days to weeks (if not longer), and it is hard to imagine an *a priori* reason that subjects with higher cognitive ability would be scanned earlier in the day, especially because they traveled to Minneapolis for the express purpose of completing this study and thus presumably were not constrained by their typical schedules. However, cognition score was not correlated with time of day for runs in session 2, and emotion score was not correlated with acquisition time for any of the six runs (Fig. S2c). Furthermore, just as for head motion, the magnitude of the correlation between time of day and cognition score did not differ across any pair of runs (all p > 0.08, Steiger’s Z test), making it unlikely that differences in prediction accuracy are influenced by time of day.

To mitigate the effect of these potential confounds, we residualized the target variable (cognition or emotion score) with respect to these two variables (head motion and time of day) before training the model. At each fold in the 10-fold cross-validation procedure, using subjects from the 9 training folds, we modeled the target variable (*y_train_*) as a linear combination of the two confounding variables and took the residual of this model *e_train_* (*y_train_ - ŷ_train_*) as input to the feature selection and model building steps. This same linear model was applied to data from subjects in the test fold to obtain a ground truth behavior score *e_test_* (*y_test_ - ŷ_test_*). To assess accuracy, model predictions were compared to *e_test_* (rather than raw score *y_test_*) as described further in the Connectome-based Predictive Modeling section below.

### Functional connectivity

For each subject and each run, we took the framewise average (at each TR) of the voxelwise signals in each of 268 nodes from the Shen atlas (Shen et al., 2013), a functional parcellation defined previously on a separate group of healthy adults. This parcellation covers the whole brain, including cortex, subcortex, and cerebellum. We then correlated all possible pairs of node timecourses to construct 268 x 268 symmetric connectivity matrices (one per subject per run). For purposes of model building, we extracted and vectorized the upper triangle of these matrices (35,778 total connections, or edges) to use as input features.

### Connectome-based Predictive Modeling

#### Overview

Connectome-based predictive modeling (CPM) is a data-driven approach that uses whole-brain functional connectivity to predict behavior (Finn et al., 2015; Shen et al., 2017). In brief, the steps involved in CPM are as follows:

1. Given a full set of subjects, each with a connectivity matrix and behavioral score, divide the data into training and test sets (here, we used 100 iterations of 10-fold cross-validation, where models were trained on 9 folds and tested on the held-out 10^th^ fold).
2. In the training set, perform mass univariate correlation between the strength of each edge (functional connectivity *r* value) and the target behavior.
3. Apply a feature selection threshold based on the magnitude and/or p-value associated with the correlation coefficients calculated in (2). This threshold is a hyperparameter that may be tuned if desired; here, we chose |r| = 0.2 (corresponding to a two-tailed p-value of approximately 0.01), since this provides good accuracy with relatively sparse features. Previous work has shown that results are generally robust to choice of threshold (Finn et al., 2015; Jangraw et al., 2018).
4. Divide edges from (3) into two tails (positive and negative) based on the sign of their correlation with behavior, then for each subject, calculate the summed strength across all edges in a given tail (X_pos_ and X_neg_).
5. Build a linear model relating the positive and negative network strengths calculated in (4) to behavior score (y):

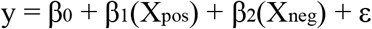
6. Calculate positive and negative network strength (X_pos_ and X_neg_) for each subject in the test set by applying the masks defined in (3) to their functional connectivity data, and use these as input to the linear model in (5) to generate predicted behavior scores.

For further details and comparisons with other predictive modeling approaches, see Shen et al. (2017).

#### Cross validation

Cross-validation was performed as follows. We first divided subjects into 10 folds respecting family structure, such that sets of siblings were always together in either the train set or the test set (never split with one sibling in the train set and another in the test set). We then trained a CPM using 9 of the 10 folds, and applied the resulting model to data from the held-out fold to generate predicted behavioral scores for all subjects in that fold. Iterating through all 10 folds yielded a vector of predicted behavioral scores for all 176 subjects in the dataset. We then repeated this entire process 100 times, to assess sensitivity of model accuracy to different fold splits.

#### Assessing model accuracy

We assessed prediction accuracy by calculating the Spearman (rank) coefficient between predicted (model generated) and observed (true) scores across subjects. Correlation is a relative measure of accuracy rather than an absolute one (e.g., mean squared error). Given that the target variables were principal component scores that were themselves made up mostly of variables measured on an arbitrary scale, we believe that relative performance—i.e., the model’s ability to distinguish higher versus lower scoring subjects—is the most appropriate metric. Because successful relative or rank prediction across subjects was our explicit goal, we used Spearman rather than Pearson correlation. However, results should be interpreted in the context of this choice, because models with good relative accuracy may still suffer from high absolute error.

#### Statistical testing

To assess the statistical significance of prediction accuracies, we generated a null distribution of expected accuracies due to chance by shuffling behavioral scores with respect to connectivity matrices and reperforming the entire analysis pipeline. A total of 10,000 randomizations were performed for each input-output combo (e.g., REST1-cognition score, MOVIE1-cognition score, inception-emotion score). We then calculated a non-parametric p-value for the observed model accuracy using the following formula,

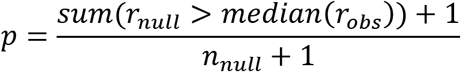

where n_null_ = 10,000 and median(r_obs_) is the median accuracy of the 100 true models.

### Anatomy of predictive networks

Due to the nature of cross-validated model building, different sets of edges may be selected at each fold of a single 10-fold cross-validation, as well as across the 100 iterations of 10-fold cross-validation. In selecting edges to visualize and interpret, for a given run, behavior score, and tail (positive and negative), we first averaged the number of times an edge was selected *within* each 10-fold run (resulting in a number between 0 and 1 for each edge), and then averaged those fractions across all 100 train-test split iterations. We then limited visualization to edges that appeared in at least 90 percent of all models, to ensure that we were considering only edges that were most robustly associated with and predictive of behavior.

We assessed significance of edge overlap across runs and videos using the hypergeometric cumulative distribution function, which gives the probability of observing a given overlap in “hits” between two binary vectors based on the total number of “hits” in each vector and the total possible hits.

For visualization purposes, we summarized edges in two ways: 1) based on individual nodes, and 2) by pooling edges into larger anatomical regions using a predefined assignment of nodes into lobes and other macroscale territories. In both cases, markers were sized by the total number of edges across both positive and negative networks (such that larger areas had more connections overall; note that individual nodes may have connections in both networks), and colored by the difference between totals in the positive and negative networks (such that red indicates more representation in the positive network, blue in the negative network, and yellow in between).

### Video clip feature extraction

We extracted video clip features in two ways: 1) by using semantic-category labels that describe high-level features of the movies made available by the HCP (*7T_movie_resources/WordNetFeatures.hdf5*) based on the approach in (Huth et al., 2012), and 2) by using the open-source package *pliers* (McNamara et al., 2017) (https://github.com/tyarkoni/pliers) to extract a number of additional features for each of the movies, including the low-level properties of brightness (luminance), vibrance, optical flow, and the mid-level property of presence of faces onscreen. Labels from (1) are provided at the same temporal resolution of the fMRI data (i.e., one set of labels per TR); we averaged these across TRs to arrive at one set of labels per video representing the average semantic content onscreen during that video. For labels in (2), we extracted features at the temporal resolution of the videos themselves (i.e., one value per frame, where the frame rate of the videos was 24 frames per second), then averaged these. In both cases, we restricted this averaging to the first 143 TRs of the videos, to match the fMRI data that were used as input to CPM.

---

**Supplemental Figure 1.**
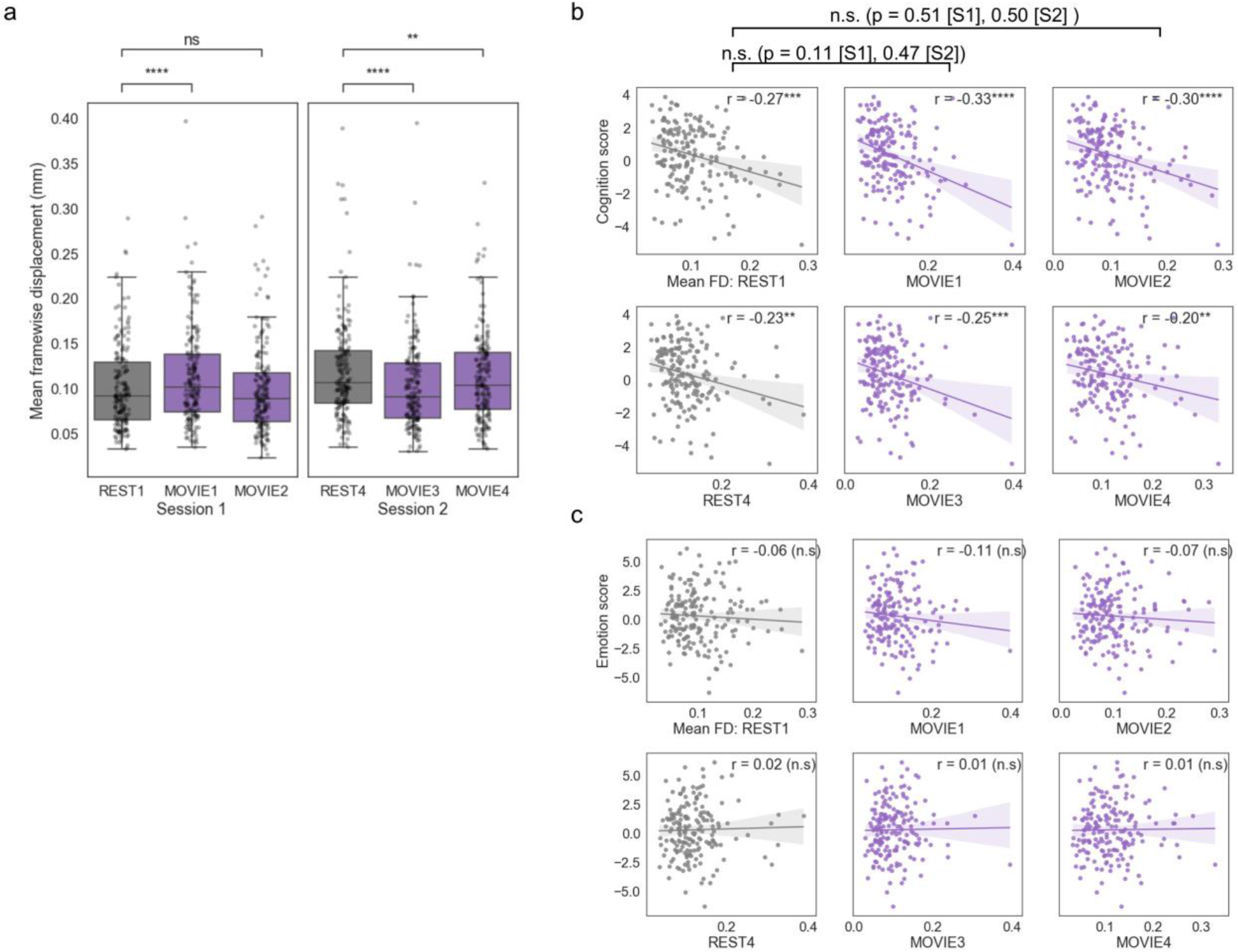
Differences in head motion across runs and correlations between head motion and behavior. a) Distributions of head motion (as indexed by mean framewise displacement [FD]) across runs. Each subject contributes one data point. b) Correlations between head motion in each run (*x* axis) and cognition score (*y* axis) across subjects. While head motion was significantly negatively associated with cognition score for all six runs, the magnitude of this correlation did not differ between runs. c) Correlations between head motion in each run (*x* axis) and emotion score (*y* axis) across subjects. Head motion was not associated with emotion score in any run. n.s., not significant (p > 0.05); *p < 0.05; **p < 0.01; ***p < 0.001; ****p < 0.0001.

**Supplemental Figure 2.**
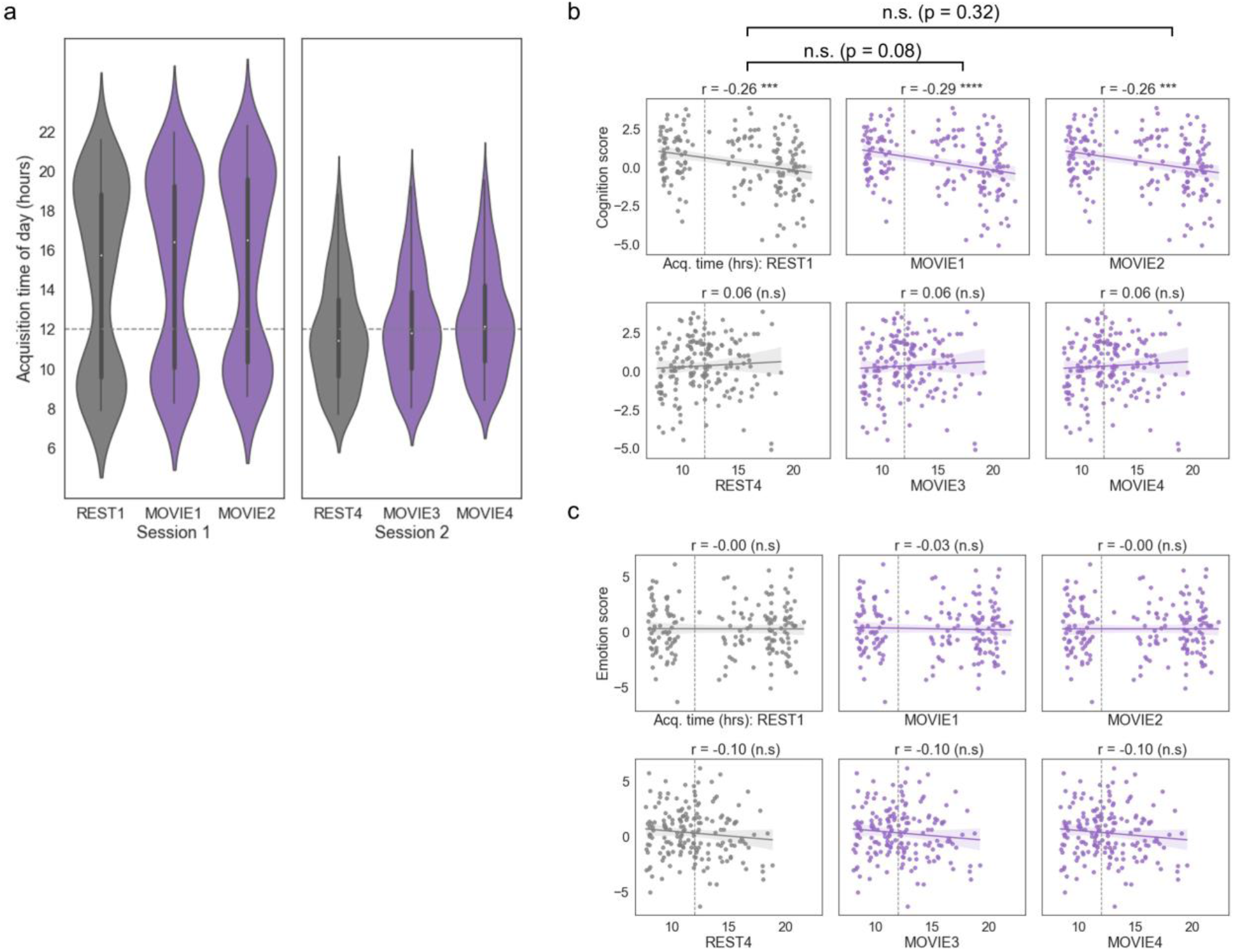
Differences in acquisition time of day across runs and correlations between time of day and behavior. a) Distributions of acquisition time of day across runs. Each subject contributes one data point. b) Correlations between acquisition time of day for each run (*x* axis) and cognition score (*y* axis) across subjects. While time of day was significantly negatively associated with cognition score across runs in session 1, the magnitude of this correlation did not differ between runs. c) Correlations between time of day of each run (*x* axis) and emotion score (*y* axis) across subjects. Time of day was not associated with emotion score in any run. In all graphs, dotted gray line represents noon (12pm). n.s., not significant (p > 0.05); *p < 0.05; **p < 0.01; ***p < 0.001; ****p < 0.0001.

**Supplemental Figure 3.**
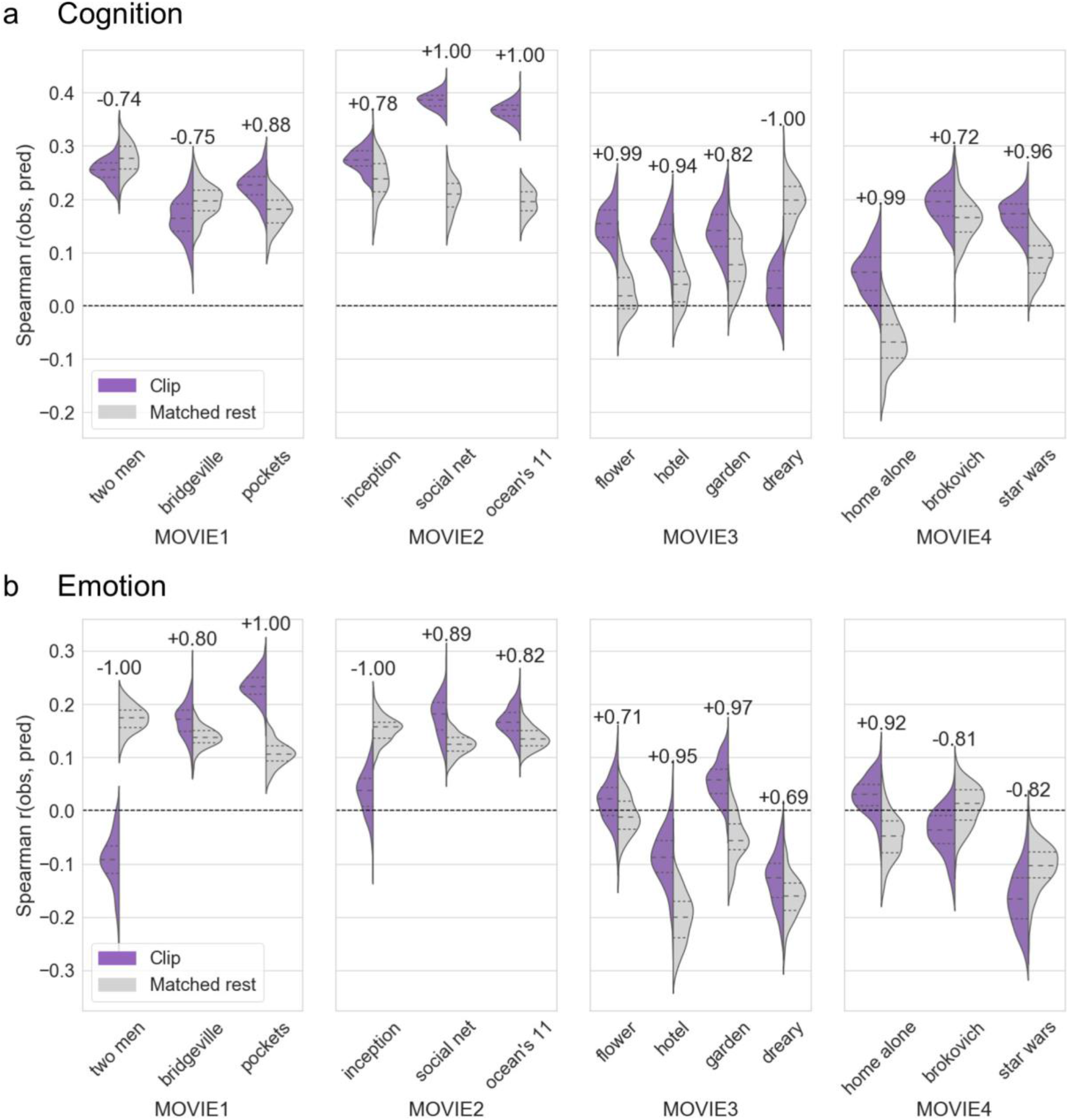
Comparing prediction accuracy between clips and their matched rest blocks. For each clip, we compared the distribution of prediction accuracies based on functional connectivity matrices derived from the clip itself to functional connectivity matrices derived from a matching set of TRs in the rest run from the corresponding session. Each state (clip or matched rest; i.e., each side of a violin) has n = 100 data points (from 100 iterations of 10-fold cross-validation). Dashed lines represent median and first and third quartiles. Clip distributions (purple) are the same as those shown in Fig. 5. Differences between states were assessed using Mann-Whitney U tests. All p-values were significant at p < 0.0001. The number written above each violin reflects the common-language effect size for the difference in accuracy between states (calculated as U/(n_1_*n_2_), or the proportion of sample pairs [1,000 total] that support a single direction). A plus sign (+) indicates that the effect is in the direction *clip > matched rest*, while a minus sign (−) indicates the opposite direction, *matched rest > clip*.

**Supplemental Figure 4.**
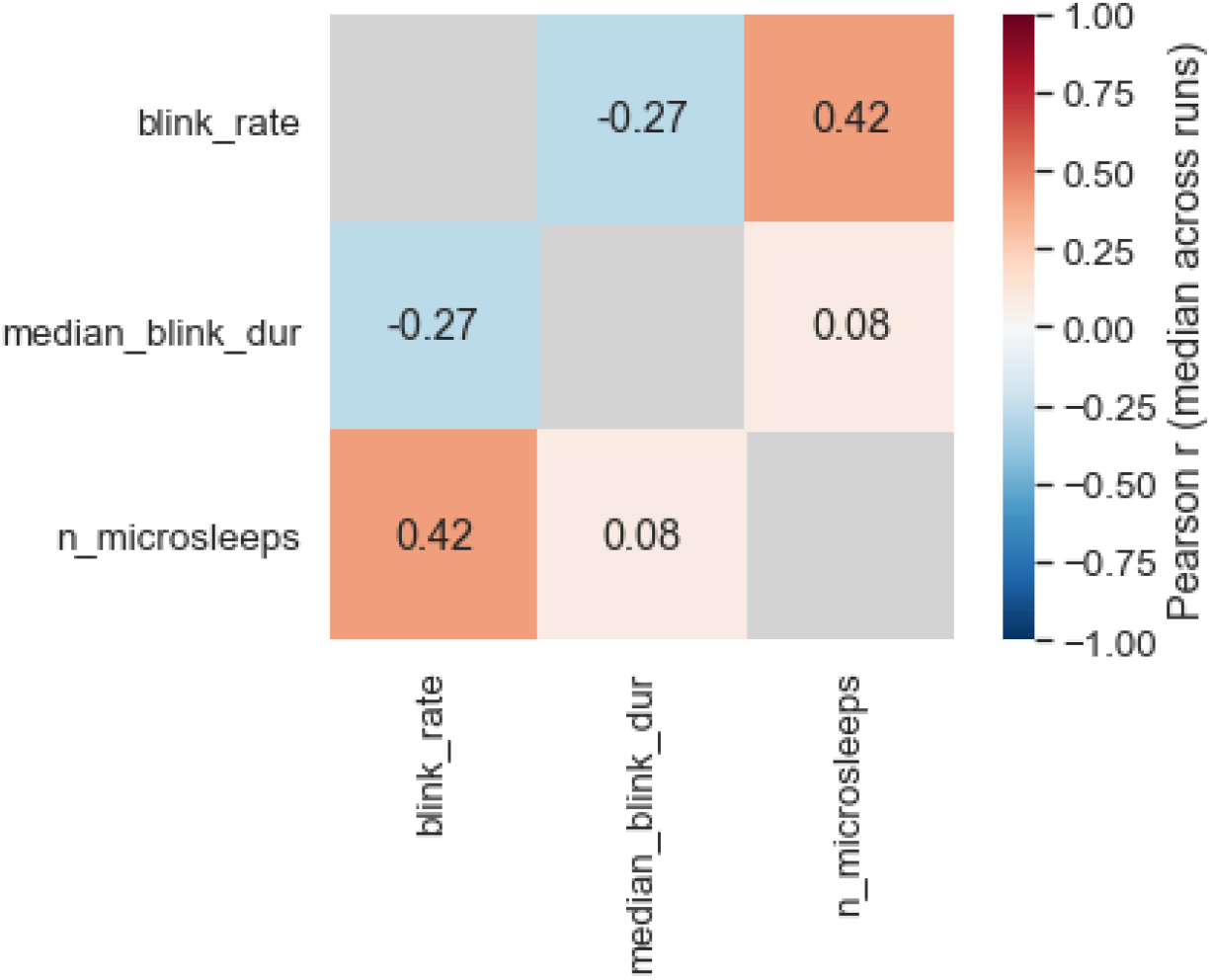
Correlations between blink-related eye tracking measures. Blink rate and duration were moderately negatively correlated, such that subjects who blinked more tended to have faster blinks. Blink rate and number of microsleeps were positively correlated, such that subjects who blinked more also tended to have more microsleeps (defined as blinks equal to or greater than 1 s in duration). Blink duration and number of microsleeps were not strongly related. Pearson r values are the median across the six runs of interest (REST1, MOVIE1, MOVIE2, REST4, MOVIE3, MOVIE4).

**Supplemental Figure 5.**
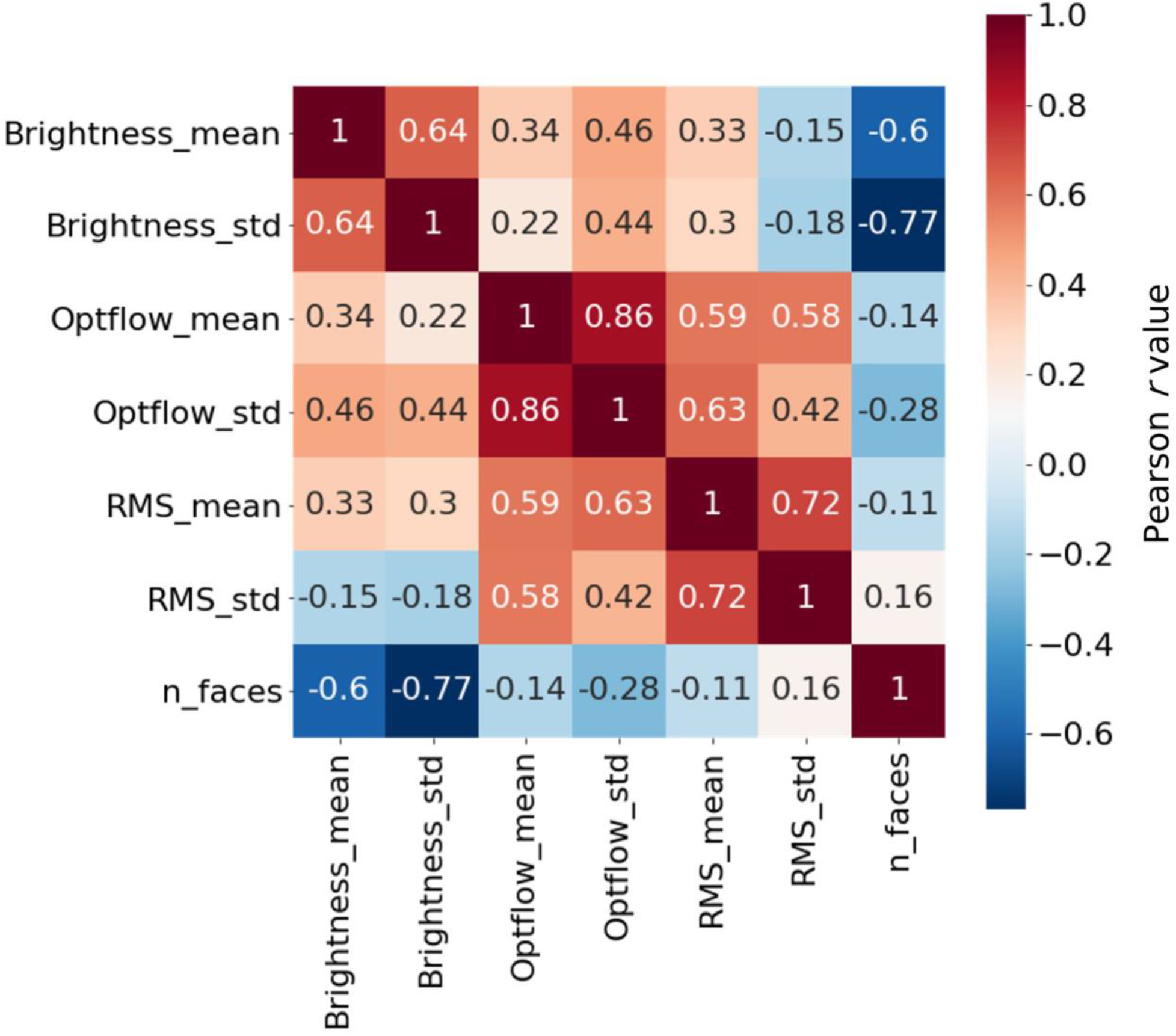
Collinearities between stimulus features. Matrix of Pearson correlation coefficients between low-level and social features extracted from individual video clips (n = 13, same clips as in Fig. 5). Optflow, optical flow; RMS, audio root-mean-square (volume); n_faces, number of TRs with at least one face onscreen. See also Fig. 7b,c.

**Table S1.**
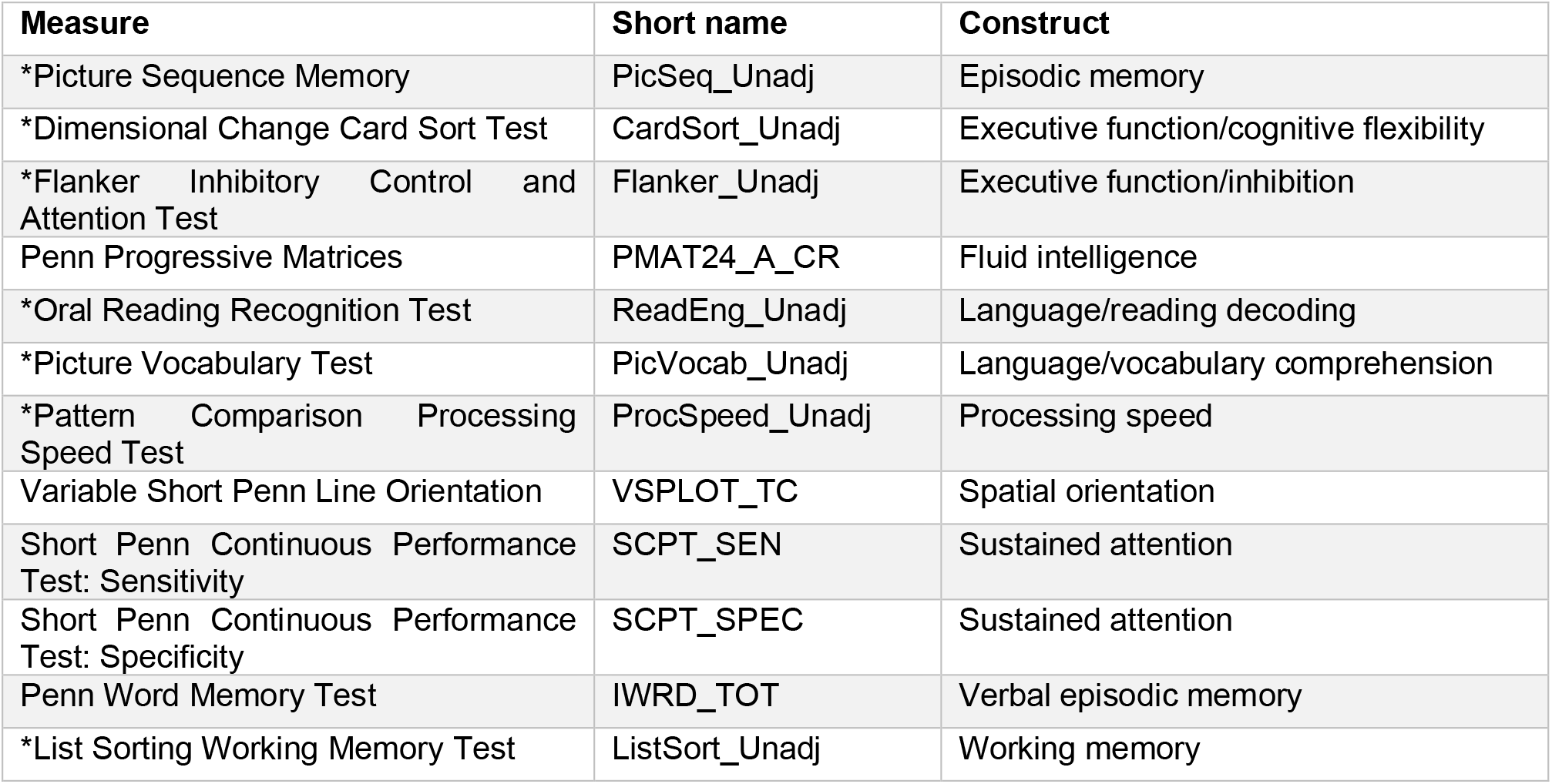
Measures from HCP Cognition domain entered into principal components analysis to derive cognition score. \**indicates item from NIH Cognition Toolbox*

**Table S2.** Measures from HCP Emotion domain entered into principal components analysis to derive emotion score. *All items from NIH Emotion Toolbox*.

| Measure | Short name | Domain |
| --- | --- | --- |
| Anger-Affect Survey | AngAffect_Unadj | Negative affect |
| Anger-Hostility Survey | AngHostil_Unadj | Negative affect |
| Anger-Physical Aggression Survey | AngAggr_Unadj | Negative affect |
| Fear-Affect Survey | FearAffect_Unadj | Negative affect |
| Fear-Somatic Arousal Survey | FearSomat_Unadj | Negative affect |
| Sadness Survey | Sadness_Unadj | Negative affect |
| General Life Satisfaction Survey | LifeSatisf_Unadj | Psychological well-being |
| Meaning and Purpose Survey | MeanPurp_Unadj | Psychological well-being |
| Positive Affect Survey | PosAffect_Unadj | Psychological well-being |
| Friendship Survey | Friendship_Unadj | Social relationships |
| Loneliness Survey | Loneliness_Unadj | Social relationships |
| Perceived Hostility Survey | PercHostil_Unadj | Social relationships |
| Perceived Rejection Survey | PercReject_Unadj | Social relationships |
| Emotional Support Survey | EmotSupp_Unadj | Social relationships |
| Instrumental Support Survey | InstruSupp_Unadj | Social relationships |
| Perceived Stress Survey | PercStress_Unadj | Stress and self-efficacy |
| Self-Efficacy Survey | SelfEff_Unadj | Stress and self-efficacy |

## REFERENCES

Bentivoglio, A.R., Bressman, S.B., Cassetta, E., Carretta, D., Tonali, P., and Albanese, A. (1997). Analysis of blink rate patterns in normal subjects. Mov Disord 12, 1028–1034.

Betti, V., Della Penna, S., de Pasquale, F., Mantini, D., Marzetti, L., Romani, Gian L., and Corbetta, M. (2013). Natural Scenes Viewing Alters the Dynamics of Functional Connectivity in the Human Brain. Neuron 79, 782–797.

Betzel, R.F., Byrge, L., Esfahlani, F.Z., and Kennedy, D.P. (2020). Temporal fluctuations in the brain’s modular architecture during movie-watching. Neuroimage 213, 116687.

Bijsterbosch, J.D., Woolrich, M.W., Glasser, M.F., Robinson, E.C., Beckmann, C.F., Van Essen, D.C., Harrison, S.J., and Smith, S.M. (2018). The relationship between spatial configuration and functional connectivity of brain regions. eLife 7, e32992.

Bolton, T.A.W., Jochaut, D., Giraud, A.L., and Ville, D.V.D. (2018). Brain dynamics in ASD during movie-watching show idiosyncratic functional integration and segregation. Hum Brain Mapp 39, 2391–2404.

Byrge, L., Dubois, J., Tyszka, J.M., Adolphs, R., and Kennedy, D.P. (2015). Idiosyncratic Brain Activation Patterns Are Associated with Poor Social Comprehension in Autism. J Neurosci 35, 5837–5850.

Chen, P.-H.A., Jolly, E., Cheong, J.H., and Chang, L.J. (2019). Inter-subject representational similarity analysis reveals individual variations in affective experience when watching erotic movies. bioRxiv, 726570.

Cole, M.W., Yarkoni, T., Repovš, G., Anticevic, A., and Braver, T.S. (2012). Global connectivity of prefrontal cortex predicts cognitive control and intelligence. J Neurosci 32, 8988–8999.

Cutting, J.E., Brunick, K.L., DeLong, J.E., Iricinschi, C., and Candan, A. (2011). Quicker, faster, darker: Changes in Hollywood film over 75 years. i-Perception 2, 569–576.

Demirtaş, M., Ponce-Alvarez, A., Gilson, M., Hagmann, P., Mantini, D., Betti, V., Romani, G.L., Friston, K., Corbetta, M., and Deco, G. (2019). Distinct modes of functional connectivity induced by movie-watching. Neuroimage 184, 335–348.

Dubois, J., and Adolphs, R. (2016). Building a Science of Individual Differences from fMRI. Trends in Cognitive Sciences 20, 425–443.

Dunbar, R.I. (1998). The social brain hypothesis. Evolutionary Anthropology: Issues, News, and Reviews: Issues, News, and Reviews 6, 178–190.

Eickhoff, S.B., Milham, M., and Vanderwal, T. (2020). Towards clinical applications of movie fMRI. Neuroimage, 116860.

Evinger, C., Manning, K.A., and Sibony, P.A. (1991). Eyelid movements. Mechanisms and normal data. Invest Ophthalmol Vis Sci 32, 387–400.

Ferguson, M.A., Anderson, J.S., and Spreng, R.N. (2017). Fluid and flexible minds: Intelligence reflects synchrony in the brain’s intrinsic network architecture. Network Neuroscience.

Finn, E.S., and Constable, R.T. (2016). Individual variation in functional brain connectivity: implications for personalized approaches to psychiatric disease. Dialogues Clin Neurosci 18, 277–287.

Finn, E.S., Corlett, P.R., Chen, G., Bandettini, P.A., and Constable, R.T. (2018). Trait paranoia shapes inter-subject synchrony in brain activity during an ambiguous social narrative. Nature Communications 9, 2043.

Finn, E.S., Glerean, E., Khojandi, A.Y., Nielson, D., Molfese, P.J., Handwerker, D.A., and Bandettini, P.A. (2020). Idiosynchrony: From shared responses to individual differences during naturalistic neuroimaging. Neuroimage 215, 116828.

Finn, E.S., Scheinost, D., Finn, D.M., Shen, X., Papademetris, X., and Constable, R.T. (2017). Can brain state be manipulated to emphasize individual differences in functional connectivity? Neuroimage 160, 140–151.

Finn, E.S., Shen, X., Scheinost, D., Rosenberg, M.D., Huang, J., Chun, M.M., Papademetris, X., and Constable, R.T. (2015). Functional connectome fingerprinting: identifying individuals using patterns of brain connectivity. Nat Neurosci 18, 1664–1671.

Gabrieli, John D.E., Ghosh, Satrajit S., and Whitfield-Gabrieli, S. (2015). Prediction as a Humanitarian and Pragmatic Contribution from Human Cognitive Neuroscience. Neuron 85, 11–26.

Geerligs, L., Rubinov, M., Cam-CAN, and Henson, R.N. (2015). State and Trait Components of Functional Connectivity: Individual Differences Vary with Mental State. J Neurosci 35, 13949–13961.

Gilson, M., Deco, G., Friston, K.J., Hagmann, P., Mantini, D., Betti, V., Romani, G.L., and Corbetta, M. (2018). Effective connectivity inferred from fMRI transition dynamics during movie viewing points to a balanced reconfiguration of cortical interactions. Neuroimage 180, 534–546.

Glasser, M.F., Sotiropoulos, S.N., Wilson, J.A., Coalson, T.S., Fischl, B., Andersson, J.L., Xu, J., Jbabdi, S., Webster, M., Polimeni, J.R., et al. (2013). The minimal preprocessing pipelines for the Human Connectome Project. Neuroimage 80, 105–124.

Gratton, C., Laumann, T.O., Nielsen, A.N., Greene, D.J., Gordon, E.M., Gilmore, A.W., Nelson, S.M., Coalson, R.S., Snyder, A.Z., Schlaggar, B.L., et al. (2018). Functional Brain Networks Are Dominated by Stable Group and Individual Factors, Not Cognitive or Daily Variation. Neuron 98, 439–452.e435.

Greene, A.S., Gao, S., Noble, S., Scheinost, D., and Constable, R.T. (2020). How tasks change whole-brain functional organization to reveal brain-phenotype relationships. bioRxiv, 870287.

Greene, A.S., Gao, S., Scheinost, D., and Constable, R.T. (2018). Task-induced brain state manipulation improves prediction of individual traits. Nature Communications 9, 2807.

Griffanti, L., Salimi-Khorshidi, G., Beckmann, C.F., Auerbach, E.J., Douaud, G., Sexton, C.E., Zsoldos, E., Ebmeier, K.P., Filippini, N., and Mackay, C.E. (2014). ICA-based artefact removal and accelerated fMRI acquisition for improved resting state network imaging. Neuroimage 95, 232–247.

Gruskin, D.C., Rosenberg, M.D., and Holmes, A.J. (2019). Relationships between depressive symptoms and brain responses during emotional movie viewing emerge in adolescence. bioRxiv, 542720.

Hampson, M., Driesen, N.R., Skudlarski, P., Gore, J.C., and Constable, R.T. (2006). Brain connectivity related to working memory performance. J Neurosci 26, 13338–13343.

Hasson, U., Avidan, G., Gelbard, H., Vallines, I., Harel, M., Minshew, N., and Behrmann, M. (2009). Shared and idiosyncratic cortical activation patterns in autism revealed under continuous real-life viewing conditions. Autism Research 2, 220–231.

Hasson, U., Nir, Y., Levy, I., Fuhrmann, G., and Malach, R. (2004). Intersubject Synchronization of Cortical Activity During Natural Vision. Science 303, 1634–1640.

Hearne, L.J., Mattingley, J.B., and Cocchi, L. (2016). Functional brain networks related to individual differences in human intelligence at rest. Sci Rep 6.

Hsu, W.-T., Rosenberg, M.D., Scheinost, D., Constable, R.T., and Chun, M.M. (2018). Restingstate functional connectivity predicts neuroticism and extraversion in novel individuals. Soc Cogn Affect Neurosci 13, 224–232.

Huijbers, W., Van Dijk, K.R.A., Boenniger, M.M., Stirnberg, R., and Breteler, M.M.B. (2017). Less head motion during MRI under task than resting-state conditions. Neuroimage 147, 111–120.

Huth, Alexander G., Nishimoto, S., Vu, An T., and Gallant, Jack L. (2012). A Continuous Semantic Space Describes the Representation of Thousands of Object and Action Categories across the Human Brain. Neuron 76, 1210–1224.

Jangraw, D.C., Gonzalez-Castillo, J., Handwerker, D.A., Ghane, M., Rosenberg, M.D., Panwar, P., and Bandettini, P.A. (2018). A functional connectivity-based neuromarker of sustained attention generalizes to predict recall in a reading task. Neuroimage 166, 99–109.

Karson, C.N. (1983). SPONTANEOUS EYE-BLINK RATES AND DOPAMINERGIC SYSTEMS. Brain 106, 643–653.

Kong, R., Li, J., Orban, C., Sabuncu, M.R., Liu, H., Schaefer, A., Sun, N., Zuo, X.-N., Holmes, A.J., and Eickhoff, S.B. (2019). Spatial topography of individual-specific cortical networks predicts human cognition, personality, and emotion. Cereb Cortex 29, 2533–2551.

Laumann, T.O., Gordon, E.M., Adeyemo, B., Snyder, A.Z., Joo, S.J., Chen, M.-Y., Gilmore, A.W., McDermott, K.B., Nelson, S.M., and Dosenbach, N.U. (2015). Functional System and Areal Organization of a Highly Sampled Individual Human Brain. Neuron.

Li, J., Kong, R., Liégeois, R., Orban, C., Tan, Y., Sun, N., Holmes, A.J., Sabuncu, M.R., Ge, T., and Yeo, B.T.T. (2019). Global signal regression strengthens association between resting-state functional connectivity and behavior. Neuroimage 196, 126–141.

McNamara, Q., De La Vega, A., and Yarkoni, T. (2017). Developing a comprehensive framework for multimodal feature extraction. Paper presented at: Proceedings of the 23rd ACM SIGKDD International Conference on Knowledge Discovery and Data Mining.

Montague, P.R., Dolan, R.J., Friston, K.J., and Dayan, P. (2012). Computational psychiatry. Trends in cognitive sciences 16, 72–80.

Nastase, S.A., Gazzola, V., Hasson, U., and Keysers, C. (2019). Measuring shared responses across subjects using intersubject correlation. Soc Cogn Affect Neurosci 14, 667–685.

Nguyen, M., Vanderwal, T., and Hasson, U. (2019). Shared understanding of narratives is correlated with shared neural responses. Neuroimage 184, 161–170.

Noble, S., Spann, M.N., Tokoglu, F., Shen, X., Constable, R.T., and Scheinost, D. (2017). Influences on the Test–Retest Reliability of Functional Connectivity MRI and its Relationship with Behavioral Utility. Cereb Cortex 27, 5415–5429.

Patzelt, E.H., Hartley, C.A., and Gershman, S.J. (2018). Computational phenotyping: using models to understand individual differences in personality, development, and mental illness. Personality Neuroscience 1.

Rosenberg, M.D., Finn, E.S., Scheinost, D., Papademetris, X., Shen, X., Constable, R.T., and Chun, M.M. (2016). A neuromarker of sustained attention from whole-brain functional connectivity. Nat Neurosci 19, 165–171.

Rosenberg, M.D., Scheinost, D., Greene, A.S., Avery, E.W., Kwon, Y.H., Finn, E.S., Ramani, R., Qiu, M., Constable, R.T., and Chun, M.M. (2020). Functional connectivity predicts changes in attention observed across minutes, days, and months. Proc Natl Acad Sci USA 117, 3797–3807.

Salehi, M., Greene, A.S., Karbasi, A., Shen, X., Scheinost, D., and Constable, R.T. (2020). There is no single functional atlas even for a single individual: Functional parcel definitions change with task. Neuroimage 208, 116366.

Salimi-Khorshidi, G., Douaud, G., Beckmann, C.F., Glasser, M.F., Griffanti, L., and Smith, S.M. (2014). Automatic denoising of functional MRI data: Combining independent component analysis and hierarchical fusion of classifiers. Neuroimage 90, 449–468.

Salmi, J., Metwaly, M., Tohka, J., Alho, K., Leppämäki, S., Tani, P., Koski, A., Vanderwal, T., and Laine, M. (2020). ADHD desynchronizes brain activity during watching a distracted multitalker conversation. Neuroimage 216, 116352.

Salmi, J., Roine, U., Glerean, E., Lahnakoski, J., Nieminen-von Wendt, T., Tani, P., Leppämäki, S., Nummenmaa, L., Jääskeläinen, I.P., Carlson, S., et al. (2013). The brains of high functioning autistic individuals do not synchronize with those of others. NeuroImage: Clinical 3, 489–497.

Shen, X., Finn, E.S., Scheinost, D., Rosenberg, M.D., Chun, M.M., Papademetris, X., and Constable, R.T. (2017). Using connectome-based predictive modeling to predict individual behavior from brain connectivity. Nat Protocols 12, 506–518.

Shen, X., Tokoglu, F., Papademetris, X., and Constable, R. (2013). Groupwise whole-brain parcellation from resting-state fMRI data for network node identification. Neuroimage 82, 403–415.

Siegel, J.S., Mitra, A., Laumann, T.O., Seitzman, B.A., Raichle, M., Corbetta, M., and Snyder, A.Z. (2016). Data quality influences observed links between functional connectivity and behavior. Cereb Cortex 27, 4492–4502.

Smith, S.M., Vidaurre, D., Beckmann, C.F., Glasser, M.F., Jenkinson, M., Miller, K.L., Nichols, T.E., Robinson, E.C., Salimi-Khorshidi, G., and Woolrich, M.W. (2013). Functional connectomics from resting-state fMRI. Trends in cognitive sciences 17, 666–682.

Sonkusare, S., Breakspear, M., and Guo, C. (2019). Naturalistic Stimuli in Neuroscience: Critically Acclaimed. Trends in Cognitive Sciences 23, 699–714.

van Baar, J.M., Chang, L.J., and Sanfey, A.G. (2019). The computational and neural substrates of moral strategies in social decision-making. Nature Communications 10, 1483.

van den Heuvel, M.P., Stam, C.J., Kahn, R.S., and Pol, H.E.H. (2009). Efficiency of functional brain networks and intellectual performance. J Neurosci 29, 7619–7624.

Van Essen, D.C., Smith, S.M., Barch, D.M., Behrens, T.E., Yacoub, E., and Ugurbil, K. (2013). The WU-Minn human connectome project: an overview. Neuroimage 80, 62–79.

Vanderwal, T., Eilbott, J., Finn, E.S., Craddock, R.C., Turnbull, A., and Castellanos, F.X. (2017). Individual differences in functional connectivity during naturalistic viewing conditions. Neuroimage 157, 521–530.

Vanderwal, T., Kelly, C., Eilbott, J., Mayes, L.C., and Castellanos, F.X. (2015). Inscapes: A movie paradigm to improve compliance in functional magnetic resonance imaging. Neuroimage 122, 222–232.

Waites, A.B., Stanislavsky, A., Abbott, D.F., and Jackson, G.D. (2005). Effect of prior cognitive state on resting state networks measured with functional connectivity. Hum Brain Mapp 24, 59–68.

Wang, J., Ren, Y., Hu, X., Nguyen, V.T., Guo, L., Han, J., and Guo, C.C. (2017). Test–retest reliability of functional connectivity networks during naturalistic fMRI paradigms. Hum Brain Mapp 38, 2226–2241.

Woo, C.-W., Chang, L.J., Lindquist, M.A., and Wager, T.D. (2017). Building better biomarkers: brain models in translational neuroimaging. Nat Neurosci 20, 365.

